# Chromatin architecture sets origin licensing capacity

**DOI:** 10.64898/2026.05.25.727720

**Authors:** Eric J. Foss, Ilana M Nodelman, Alexa Usenko, Holden Furutani, Ayush Goel, Brandon Lofts, Shawna Miles, Zhiguo Zhang, Gregory D Bowman, Toshio Tsukiyama, Antonio Bedalov

**Affiliations:** Division of Translational Sciences and Therapeutics and Human Biology, Fred Hutchinson Cancer Center, Seattle, WA, 98109, USA; Division of Basic Sciences, Fred Hutchinson Cancer Center, Seattle, WA, 98109, USA; Thomas C. Jenkins Department of Biophysics, Johns Hopkins University, Baltimore, MD, 21218, USA; Institute for Cancer Genetics, Department of Pediatrics and Department of Genetics and Development, Columbia University Irving Medical Center, New York, NY, 10032, USA

## Abstract

Replication origin licensing enables complete and faithful genome duplication, yet how chromatin regulates this process *in vivo* remains unclear. Using MCM–ChEC–seq to track helicase loading from metaphase through G1 in budding yeast, we find that licensing occurs in a rapid, synchronous burst at mitotic exit and then reaches an early plateau despite continued permissive cell-cycle conditions and persistent ORC binding at origins. Here we show that this plateau is imposed by chromatin architecture at replication origins, which limits the extent of origin licensing. Histone H3K56 acetylation marks newly replicated chromatin and is removed at S-phase exit by the deacetylases Hst3 and Hst4. Persistent H3K56ac severely impairs MCM loading without affecting ORC occupancy, indicating that chromatin limits licensing at the helicase-loading step. Strikingly, deletion or catalytic inactivation of the chromatin remodeler Isw2 increases licensing by approximately 40% in wild-type cells and fully suppresses the licensing defect in *hst3*Δ *hst4*Δ mutants, identifying Isw2 as a physiological inhibitor of origin licensing. Isw2-dependent nucleosome repositioning narrows the origin nucleosome-depleted region and restricts helicase loading. Together, these findings show that chromatin architecture at replication origins sets licensing capacity. Newly replicated chromatin transiently adopts an Isw2-dependent inhibitory configuration that is relieved, but not completely eliminated, by post-replicative chromatin maturation. Genome-wide licensing thus reflects integration of chromatin-imposed licensing capacity with cell cycle-dependent control of licensing timing.

## Introduction

Faithful duplication of the eukaryotic genome requires that replication origins initiate once and only once per cell cycle (Diffley 1996). This control is enforced by temporally separating origin licensing from origin firing through cyclin-dependent kinase (CDK) activity (Blow and Dutta 2005). During late mitosis and G1, when CDK activity is low, origins become licensed through ordered assembly of pre-replicative complex components, culminating in loading of the Mcm2–7 helicase as a head-to-head double hexamer (Remus et al. 2009; Gambus et al. 2011). Upon entry into S phase, rising CDK activity both activates licensed origins and simultaneously inhibits new licensing events (Tanaka and Diffley 2002), thereby enforcing once-per-cycle replication. While this regulatory framework defines when licensing is permitted, it does not address whether replication origins *in vivo* are intrinsically competent to support helicase loading.

The molecular requirements for origin licensing are well defined biochemically. The origin recognition complex (ORC) binds replication origins and, together with Cdc6 and Cdt1, promotes MCM loading through ordered conformational transitions that are inhibited by CDK activity (Bell and Stillman 1992; Tanaka and Diffley 2002; Remus et al. 2009). Biochemical, structural, and single-molecule studies show that helicase loading requires sequential loading of two oppositely oriented MCM hexamers by a single ORC complex, which must reposition and reorient between loading reactions (Ticau et al. 2015; Coster and Diffley 2017; Gupta et al. 2021). This reaction therefore requires accessible DNA flanking the origin to accommodate double-hexamer assembly. In cells, however, origin DNA resides within chromatin. In budding yeast, origins occupy compact nucleosome-depleted regions (NDR), where loading of the MCM double hexamer sterically occludes the ORC-binding site and promotes ORC dissociation from licensed origins (Reuter et al. 2024; Speck and Reuter 2025). Given this confined architecture, nucleosome positioning near origins could constrain helicase loading even when ORC binding is intact.

In cells, origin licensing is restricted to a defined interval of low CDK activity extending from mitotic exit through G1. Early studies documented the appearance of prereplicative complexes at mitotic exit and their persistence throughout G1, establishing this interval as permissive for licensing (Diffley 1994; Dahmann et al. 1995; Piatti et al. 1996). Because G1 lies entirely within this low-CDK window, licensing is generally assumed to accumulate progressively throughout G1 until the cell cycle advances or licensing factors become limiting. However, genome-wide measurements of licensing kinetics *in vivo* have been lacking. An alternative possibility is that helicase loading is constrained not only by cell-cycle regulation but also by chromatin architecture, such that origins are licensed rapidly upon entry into the low-CDK window but only to the extent permitted by their local chromatin environment. In this scenario, chromatin structure would set an intrinsic upper bound on licensing efficiency *in vivo*.

A likely source of such a capacity limit is chromatin organization at replication origins. In budding yeast, origins reside within NDRs flanked by positioned nucleosomes shaped by intrinsic DNA sequence features, ORC binding, and ATP-dependent remodeling activities (Simpson 1990; Lipford and Bell 2001; Berbenetz et al. 2010; Eaton et al. 2010). Because loading of the Mcm2–7 double hexamer requires free DNA flanking the origin, nucleosome positioning near origins is likely to influence licensing capacity *in vivo*.

Chromatin at replication origins is further shaped by replication itself. Passage of the replication fork disrupts nucleosome organization and is followed by rapid chromatin reassembly (Annunziato 2005; Groth et al. 2007). Newly synthesized histones are deposited in a hyperacetylated state marked by acetylation of histone H3 on lysine 56 (H3K56ac) (Masumoto et al. 2005; Xu et al. 2005). This modification, catalyzed by Rtt109, is associated with increased nucleosome mobility and enhanced chromatin remodeling (Han et al. 2007; Kaplan et al. 2008; Duan et al. 2025). At the end of S phase, H3K56ac is removed by the Hst3 and Hst4 deacetylases, stabilizing nucleosome positioning and restoring mature chromatin structure (Celic et al. 2006; Duan et al. 2025). Mutants defective in post-replicative chromatin maturation exhibit replication defects and DNA damage sensitivity (Masumoto et al 2005), but the molecular basis of these phenotypes remains unclear. These phenotypes have been attributed to impaired fork stability, changes in checkpoint activation, or delayed origin activation (Celic et al., 2008; Maas et al., 2006; Thaminy et al., 2007; Simoneau et al. 2015; Tremblay et al., 2023). However, whether chromatin maturation directly regulates helicase loading during origin licensing in the subsequent cell cycle has not been determined.

ATP-dependent chromatin remodelers provide a potential mechanistic link between post-replicative chromatin maturation and origin licensing. Remodelers including Isw1, Isw2, Ino80, and Chd1 can, together with ORC, generate phased nucleosome arrays flanking replication origins *in vitro* (Azmi et al. 2017; Chacin et al. 2023). These activities are generally viewed as facilitating origin function and accessibility. However, biochemical reconstitution experiments reveal that individual remodelers impose distinct nucleosome spacing and NDR geometries, indicating that not all chromatin configurations are equally permissive for MCM loading (Azmi et al. 2017; Chacin et al. 2023). Whether specific remodeling activities actively restrain helicase loading *in vivo* remains unknown.

These considerations raise three questions: when within the low-CDK interval origin licensing occurs *in vivo*; whether licensing is limited primarily by cell-cycle regulation or by chromatin architecture; and whether individual chromatin remodelers can restrict helicase loading despite their apparent redundancy.

Here, we directly measure genome-wide helicase loading during the M-to-G1 transition in budding yeast. Origin licensing occurs in a rapid, synchronous burst at mitotic exit rather than accumulating throughout G1, indicating that licensing competence is already established at the time of CDK inactivation. However, licensing does not proceed to completion: MCM loading stops even while CDK activity remains low and ORC remains associated with origins. We show that this limitation arises from chromatin at the origin. The remodeler Isw2 restricts helicase loading, and completion of post-replicative chromatin maturation relieves—but does not eliminate—this constraint. These findings demonstrate that chromatin architecture limits licensing capacity and that chromatin assembled during the preceding S phase encodes licensing potential for the next cell cycle.

## Results

### Rapid and synchronous burst of origin licensing at mitotic exit

To determine when origin licensing occurs during the cell cycle, we monitored MCM loading using MCM–ChEC (chromatin endogenous cleavage), in which micrococcal nuclease (MNase) fused to an MCM subunit cleaves DNA adjacent to loaded helicase complexes upon calcium addition (Figure S1). Cleavage events can be detected genome-wide by sequencing (ChEC–seq) or at specific origins by qPCR (ChEC–qPCR). Because cleavage occurs only when an MCM double hexamer is loaded, ChEC–qPCR measures the fraction of origins that are licensed in the cell population. The quantitative relationship between ChEC–qPCR signal and licensed origin fraction was previously validated (Foss et al., 2021).

Using MCM–ChEC, we monitored helicase loading during synchronous cell-cycle progression in budding yeast cells released from a G2/M arrest while following cell-cycle progression by flow cytometry. Signals were normalized using *S. pombe* spike-in reads to control for variation in MNase cleavage efficiency, DNA recovery, and sequencing depth (see Methods).

Normalized genome-wide MCM–ChEC–seq signal across 326 annotated replication origins became detectable within 20–30 minutes after release and increased sharply over the following ∼20 minutes (Figure 1A; replicate experiment in Figure S2). In parallel, ChEC–qPCR measurements at representative origins (ARS1103 and ARS224) allowed quantification of the fraction of licensed origins in the population. By 40 minutes, when only ∼6% of cells had entered G1, ∼37% of origins were already licensed, and licensing approached a plateau of ∼70–75% by 60 minutes (Figure 1B, upper panel).

**Figure 1.**
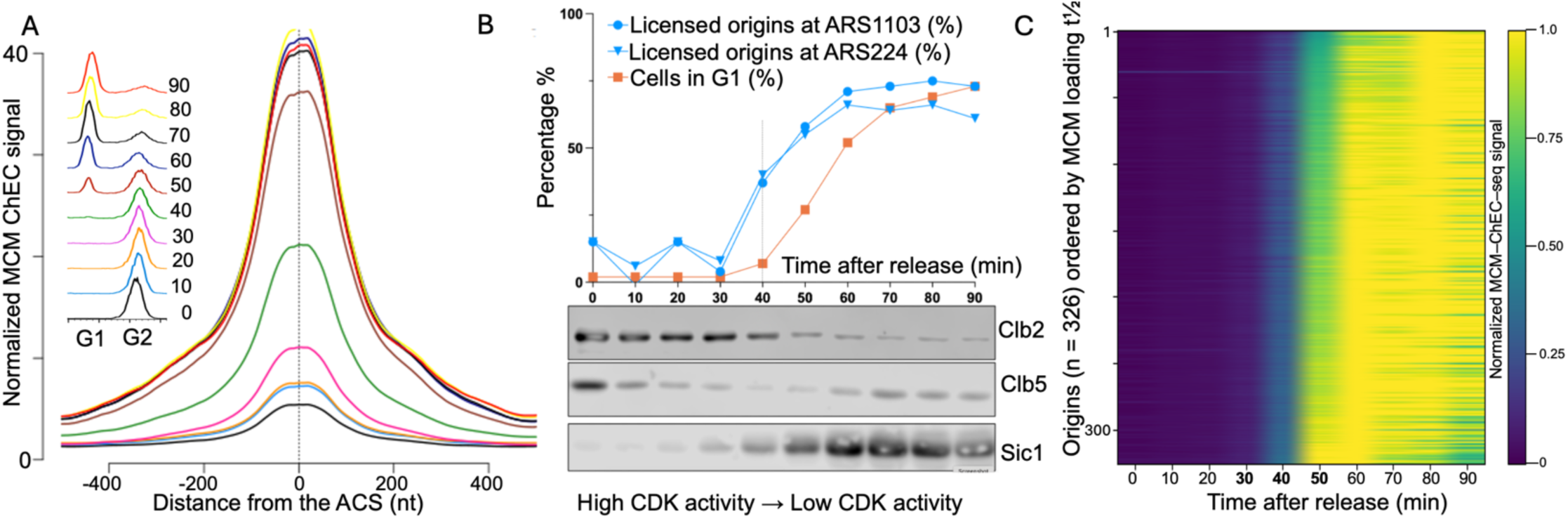
Replication origin licensing occurs rapidly and synchronously at mitotic exit. (A) Aggregate MCM loading profile after release from a nocodazole arrest. Normalized MCM–ChEC–seq signal averaged across 326 annotated replication origins and aligned relative to the ACS is shown for the indicated times after release. MCM signal is low in G2/M and increases sharply within ∼20–40 min after release, reaching near-maximal levels by ∼50–60 min. (B) Quantification of licensing and CDK regulatory markers during the same time course. The fraction of licensed origins at two representative origins (ARS1103 and ARS224) was measured by ChEC–qPCR (blue) and compared with the fraction of cells in G1 determined by flow cytometry (orange). Licensing is detectable before the majority of cells have entered G1 and reaches near-maximal levels by ∼60 min. Immunoblot analysis of Clb2 and Clb5 together with the CDK inhibitor Sic1 shows loss of B-type cyclins and accumulation of Sic1 coincident with the onset of MCM loading, consistent with licensing occurring as CDK activity declines. qPCR results are available in Supplemental Table S5. (C) Licensing kinetics at individual origins. Heatmap of normalized MCM–ChEC signal across 326 annotated replication origins, aligned relative the ACS as in (A). Origins are ordered by the time to half-maximal MCM loading (t½). Most origins acquire MCM signal within the same narrow time window, producing a near-synchronous increase in genome-wide helicase loading rather than progressive origin-by-origin licensing. MCM abundance and t½ values for individual origins are provided in Supplemental Table S4.

Thus, most origin licensing occurs shortly after release from G2/M arrest, before the majority of cells have entered G1, indicating that licensing occurs in a rapid and synchronous burst at mitotic exit rather than accumulating gradually throughout G1.

Because origin licensing is inhibited by B-type cyclin–associated CDK activity, we examined the levels of the B-type cyclins Clb2 and Clb5 together with the CDK inhibitor Sic1 across the same time course. The onset of MCM loading coincided with rapid loss of both cyclins and the concomitant accumulation of Sic1, a combination expected to extinguish Clb–CDK activity (Schwob et al., 1994) (Figure 1B, lower panel). These changes indicate that helicase loading accompanies CDK inactivation at mitotic exit.

Analysis at individual origins revealed remarkably uniform kinetics across the genome. Strikingly, the time to half-maximal MCM loading (t½) across the origin population spans only ∼10 minutes, indicating that genome-wide helicase loading occurs within a remarkably narrow temporal window. Heatmap visualization of normalized MCM–ChEC signal across 326 origins showed a near-simultaneous increase in loading rather than progressive origin-by-origin accumulation (Figure 1C). Most origins increased sharply over the same narrow time window, producing an abrupt vertical transition in the heatmap. Consistent with this observation, 307 of 326 origins (96%) reached ≥50% of maximal signal by 50 minutes after release. Licensing therefore occurs as a coordinated genome-wide transition at mitotic exit rather than accumulating progressively throughout G1.

Despite this global synchrony, reproducible differences in origin-specific licensing kinetics were evident. Comparison of the time to half-maximal MCM loading between two independent experiments showed strong correlation (Pearson r = 0.88), indicating that these origin-specific differences are highly reproducible (Figure S2D).

We therefore asked whether major origin classes exhibit systematic differences in licensing kinetics. Origins previously classified as early- or late-replicating showed largely overlapping distributions of the time to half-maximal MCM loading in both experiments (Figure S3). Thus, although individual origins differ modestly in licensing kinetics, these differences do not segregate into early- and late-replicating classes, indicating that the determinants of replication timing are not established at the licensing step.

An independent replicate experiment reproduced the aggregate kinetics and genome-wide heatmap structure (Figure S2), confirming that the synchronous burst of origin licensing at mitotic exit is a robust and reproducible feature of the system.

Because helicase loading begins immediately upon CDK inactivation, origins must already be competent for licensing at mitotic exit. This behavior suggests that the determinants of licensing competence are established prior to mitotic exit. We therefore hypothesized that post-replicative chromatin maturation at S-phase exit establishes the licensing competence that becomes revealed upon CDK inactivation.

### Isw2-dependent chromatin remodeling limits replication origin licensing

To determine whether post-replicative chromatin maturation contributes to licensing competence, we measured MCM loading in mutants affecting histone H3K56 acetylation. Using ChEC–qPCR at ten replication origins, we found that *rtt109*Δ cells exhibited a modest reduction in MCM loading, whereas *hst3*Δ *hst4*Δ cells showed a pronounced licensing defect, with a mean decrease of 38% (range 18–71%) (Figure 2A; Figure S4). These results indicate that persistent H3K56 acetylation strongly impairs origin licensing. A previous study reported reduced replication initiation in *hst3*Δ *hst4*Δ cells (Tremblay et al., 2023), suggesting that the licensing defect observed here may underlie this phenotype.

**Figure 2.**
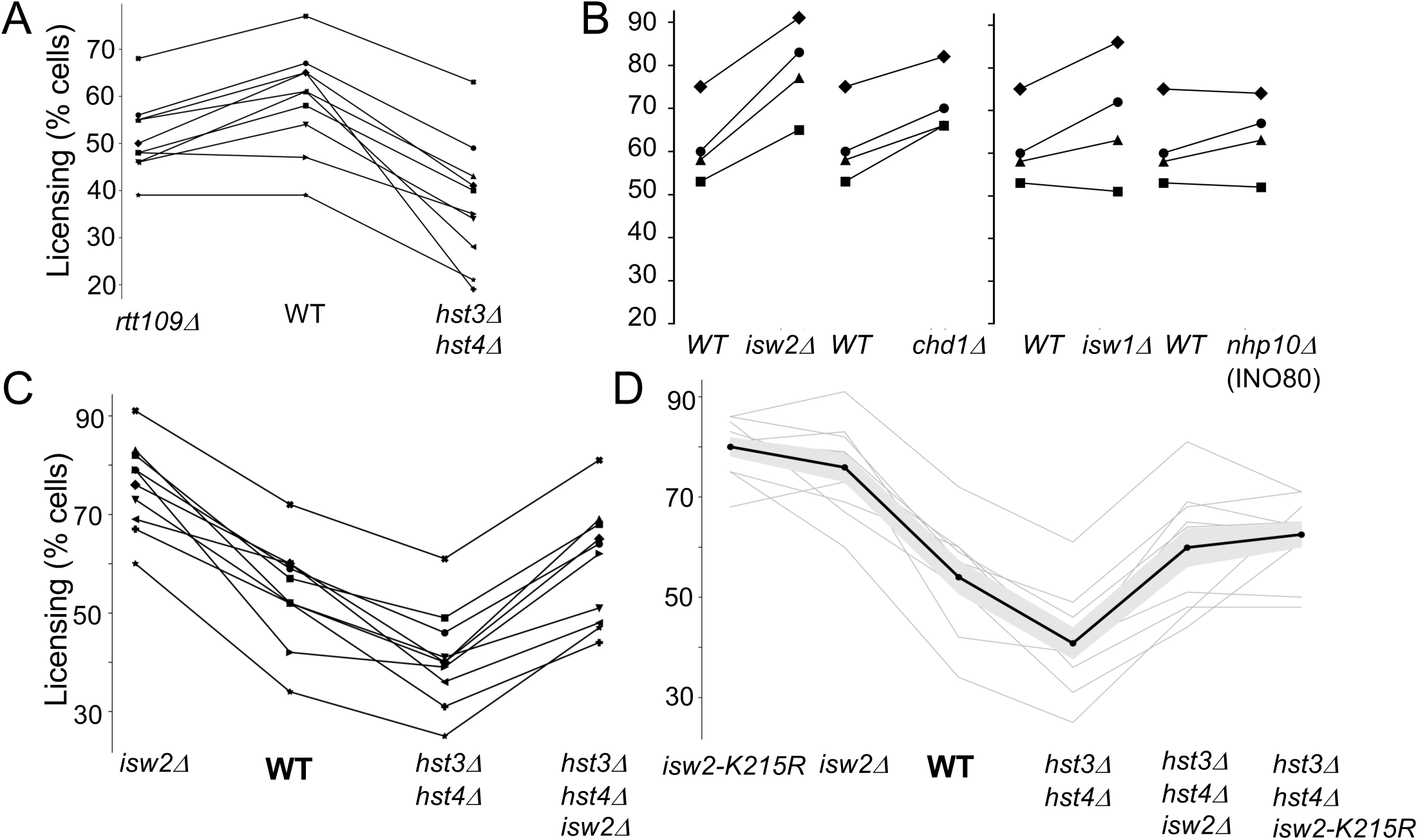
Isw2-dependent chromatin remodeling limits replication origin licensing. **(A)** MCM loading measured by ChEC–qPCR at ten replication origins (ARS216, ARS224, ARS315, ARS419, ARS605, ARS802, ARS1103, ARS1235, ARS1406, and ARS1427) in the indicated strains. Loss of H3K56 acetylation in *rtt109*Δ cells caused a modest but reproducible reduction in licensing relative to wild type (mean −13%; range −25% to +2%; paired two-tailed t-test p = 9.1 × 10⁻⁴). In contrast, persistent H3K56 acetylation in *hst3*Δ *hst4*Δ cells produced a pronounced licensing defect (mean −38%; range −71% to −18%; p = 7.4 × 10⁻⁵). Individual qPCR values are provided in Supplemental Table S7. **(B)** MCM loading in wild-type cells lacking individual chromatin remodelers. Deletion of *ISW2* significantly increased licensing (mean +29%; range +21% to +38%; p = 4.9 × 10⁻³), and deletion of *CHD1* produced a smaller increase (mean +16%; range +9% to +25%; p = 5.6 × 10⁻³). Deletion of *ISW1* or *NHP10* (Ino80 complex) did not significantly alter licensing (p = 0.14 and p = 0.31, respectively). Individual qPCR values are provided in Supplemental Table S9. **(C)** Licensing measured at ten replication origins in wild-type, *isw2*Δ, *hst3*Δ *hst4*Δ, and *isw2*Δ *hst3*Δ *hst4*Δ strains. Deletion of *ISW2* increased licensing at all origins (mean +44%; range +15% to +88%; p = 1.4 × 10⁻⁵). The licensing defect in *hst3*Δ *hst4*Δ cells (mean −24%; range −40% to −7%; p = 1.3 × 10⁻⁴ versus WT) was strongly suppressed by deletion of *ISW2* (mean +49% relative to *hst3*Δ *hst4*Δ; range +24% to +88%; p = 3.6 × 10⁻⁶). **(D)** An ATPase-dead allele of *ISW2* (*isw2*-K215R) phenocopied *isw2*Δ in both wild-type and *hst3*Δ *hst4*Δ backgrounds. Differences between deletion and catalytic mutant alleles were not significant (p = 0.13 and p = 0.41, respectively), indicating that catalytic remodeling activity is required for inhibition of origin licensing. In (D), thin gray lines represent individual origins; the thick black line indicates the mean across origins, and the shaded region denotes ± SEM. Individual qPCR values are provided in Supplemental Table S6. All statistical comparisons were performed using paired two-tailed t-tests across origins.

Because helicase loading requires accessible DNA adjacent to replication origins, we next examined ATP-dependent chromatin remodelers previously implicated in phasing origin-flanking nucleosomes *in vitro* (Isw1, Isw2, Ino80, and Chd1) (Chacin et al. 2023). Measuring MCM loading in otherwise wild-type cells lacking individual remodeler activities revealed remodeler-specific effects on licensing. Deletion of *ISW2* produced a large and consistent increase in MCM loading at all origins examined (Figure 2B), whereas deletion of *CHD1* caused a more modest increase. In contrast, deletion of *ISW1* or *NHP10*—which encodes an accessory subunit of the INO80 complex—had little detectable effect on licensing under these conditions (Figure 2B).

We next examined *ISW2* deletion at six additional replication origins in both wild-type and *hst3*Δ *hst*4Δ backgrounds, extending the analysis to ten origins. In wild-type cells, *ISW2* deletion increased licensing at every origin (average 44%, range 17–88%) (Figure 2C), indicating that Isw2 normally restrains MCM loading under conditions of proper chromatin maturation. In *hst3*Δ *hst4*Δ cells, *isw2*Δ produced an even larger increase (58%), effectively suppressing the licensing defect caused by persistent H3K56 acetylation. At most origins (8 of 10), licensing in *isw2*Δ *hst3*Δ *hst4*Δ cells exceeded wild-type levels. An ATPase-dead Isw2 allele fully phenocopied *isw2*Δ in both wild-type and *hst3*Δ *hst4*Δ backgrounds (Figure 2D), demonstrating that catalytic remodeling activity is required for this inhibitory effect. An independent ChEC–qPCR experiment reproduced these results across the same set of origins (Figure S5).

Consistent with these targeted measurements, genome-wide MCM ChIP–seq—an orthogonal assay—showed reduced origin-associated MCM signal in *hst3Δ hst4Δ* and increased signal in *isw2Δ*, with strong suppression of the *hst3Δ hst4Δ* licensing defect by *ISW2* deletion (Figure S6). These results indicate that the licensing phenotypes extend genome-wide across replication origins.

Together, these results identify Isw2 as a chromatin remodeler that restrains origin licensing under normal conditions, as loss of *ISW2* increased MCM loading beyond wild-type levels at all tested origins. This effect could reflect reduced recruitment of the ORC to replication origins or inhibition of the downstream helicase-loading reaction after ORC recruitment. To distinguish between these possibilities, we next examined ORC occupancy.

### Isw2 limits origin licensing after ORC recruitment

To determine whether Isw2 restrains origin licensing by affecting ORC recruitment or a downstream step in helicase loading, we measured ORC occupancy at replication origins by ChIP in G2/M and G1 cells from WT, *hst3*Δ *hst4*Δ, and *isw2*Δ strains. To enable spatial interpretation of helicase loading, we analyzed a curated set of 91 replication origins with precisely mapped ORC position and polarity based on ORC–ChEC-seq and ORC ChIP-exo datasets (Chappleboim et al. 2024; Reuter et al. 2024). Across these oriented origins, the ORC ChIP peak lies to the left of the MCM midpoint used for alignment, confirming the expected spatial relationship between ORC binding and the site of MCM loading and validating the orientation of the curated origin set (Figure 3).

**Figure 3.**
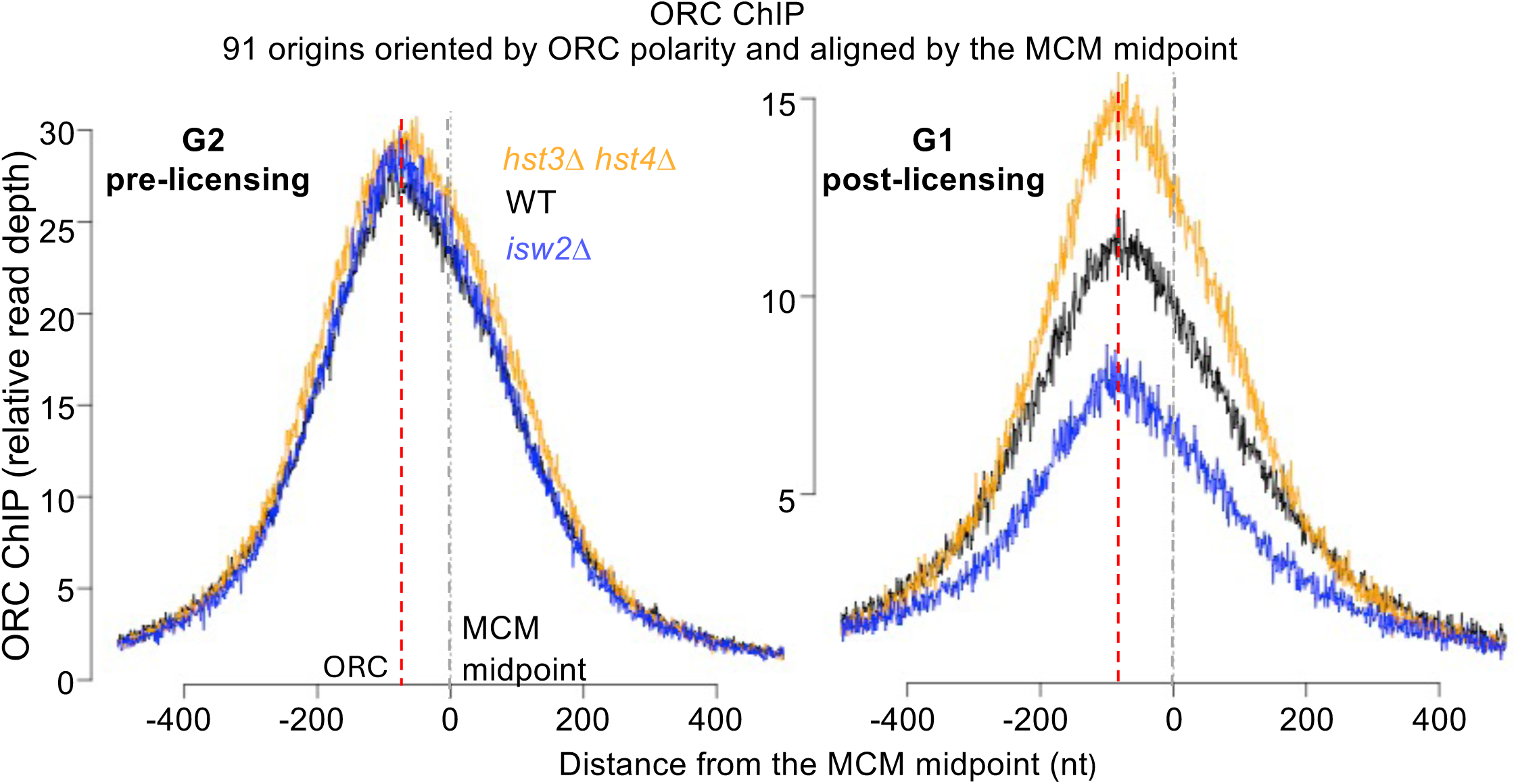
ORC recruitment is unchanged across genotypes, but post-licensing ORC retention differs. ORC ChIP signal was measured at a curated set of 91 replication origins in synchronized WT, *hst3*Δ *hst4*Δ, and *isw2*Δ cells and plotted relative to the MCM loading midpoint (0 bp). Left, G2/M (pre-licensing): ORC occupancy profiles were indistinguishable among genotypes, indicating that altered chromatin maturation and loss of Isw2 do not affect ORC recruitment to origins. Right, G1 (post-licensing): ORC occupancy diverged, with highest retention in *hst3*Δ *hst4*Δ, intermediate levels in wild type, and lowest levels in *isw2*Δ. Because stable ORC association decreases upon successful MCM double-hexamer assembly, G1 ORC retention inversely reports productive licensing. The persistence of substantial ORC signal at wild-type origins indicates that a subset of origins remains unlicensed under licensing-permissive conditions, and that chromatin regulators modulate the extent of helicase loading without altering initial ORC recruitment. Relative ORC ChIP signal at each of the 91 origins is provided in Supplemental Table S10.

In G2/M, ORC occupancy was comparable across all strains, indicating that chromatin maturation and Isw2 do not affect ORC recruitment. In G1, however, ORC occupancy diverged among strains, being highest in *hst3*Δ *hst4*Δ, intermediate in WT, and lowest in *isw2*Δ—the inverse of the licensing measured at individual origins by ChEC–qPCR (Figure 2) and genome-wide by MCM ChIP (Figure S6). Because stable ORC association decreases after successful MCM double-hexamer assembly (Reuter et al. 2024), G1 ORC occupancy is expected to inversely report the fraction of origins that remain unlicensed, as observed here. Importantly, ORC remained detectable at wild-type origins under licensing-permissive G1 conditions, indicating that a subset of ORC-bound origins remains unlicensed even when CDK activity is low. This fraction increased in *hst3Δ hst4Δ* and decreased in *isw2*Δ. Thus, permissive CDK activity and origin-bound ORC are not sufficient to ensure helicase loading. This graded response indicates that Isw2-dependent chromatin structure limits helicase loading downstream of ORC recruitment.

Similar ORC occupancy patterns were observed in an independent ChIP experiment in which chromatin was fragmented using exogenous MNase (Figure S7).

MCM loading requires engagement of DNA adjacent to the ACS, suggesting that a physical constraint at the loading site limits helicase assembly. Because the curated origin set has experimentally confirmed ORC polarity and a defined direction of MCM loading, chromatin organization can be interpreted relative to the site of helicase assembly. We therefore asked whether Isw2 regulates origin nucleosome organization in a manner that modulates helicase loading.

### Isw2-dependent nucleosome positioning narrows the origin NDR

To examine origin chromatin structure independently of cellular processes, we analyzed published biochemical reconstructions of chromatin assembled on purified origin DNA together with ORC and individual remodelers (Chacin et al. 2023). Because these reactions occur in the absence of transcription, replication, and other nuclear activities, nucleosome positioning reflects the intrinsic action of remodelers on origin DNA.

ORC together with individual chromatin remodelers (Isw1, Isw2, Ino80, and Chd1) generates phased nucleosome arrays flanking replication origins, whereas neither ORC alone nor remodelers acting in isolation are sufficient to establish this organization (Chacin et al. 2023). Because nucleosome positions are poorly defined in the absence of ORC- and remodeler-dependent phasing, we compared the ORC-phased origin chromatin architecture generated by Isw2 with that produced by Isw1, Ino80, or Chd1 across the same curated set of 91 oriented origins, enabling remodeler-specific differences in nucleosome positioning to be interpreted relative to the helicase-loading reaction. This analysis recapitulated remodeler-specific differences in origin architecture: Isw2, and to a lesser extent Chd1, shifted origin-flanking nucleosomes inward relative to Isw1 or Ino80, thereby narrowing the nucleosome-depleted region (NDR) at these origins (Figure 4).

**Figure 4.**
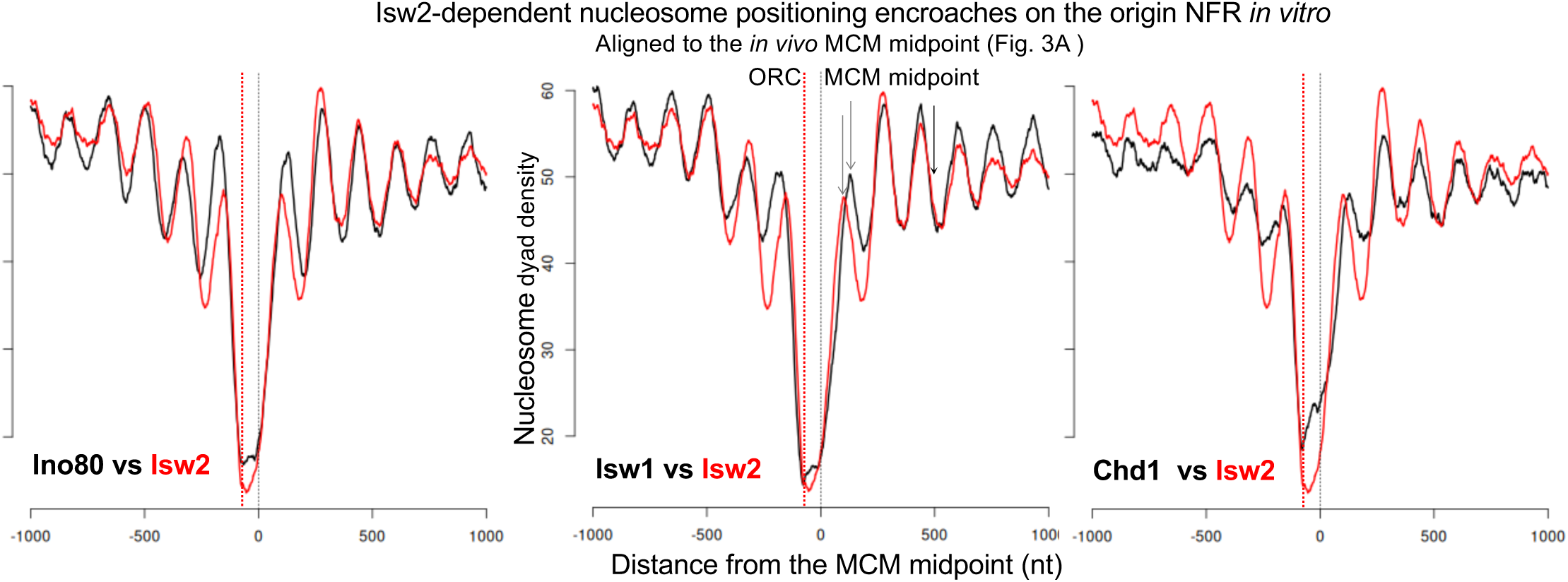
Isw2-dependent nucleosome positioning narrows the origin NDR relative to other remodelers *in vitro*. Published biochemical reconstructions of chromatin assembled on purified origin DNA together with ORC and individual chromatin remodelers Chacin et al. 2023 were reanalyzed using the same curated set of 91 oriented replication origins examined *in vivo*. Nucleosome dyad density profiles were aligned to the *in vivo* MCM midpoint, as in Figure 3, allowing origin chromatin organization to be interpreted relative to the helicase-loading reaction. Vertical dashed lines indicate the ORC binding site and the MCM midpoint. Direct comparison of ORC-phased chromatin remodeled by Isw2 with that remodeled by Isw1, Ino80, or Chd1 reveals remodeler-specific differences in nucleosome positioning. Relative to Isw1 and Ino80, Isw2 promotes inward positioning of origin-flanking nucleosomes, thereby narrowing the nucleosome-depleted region (NDR), whereas Chd1 produces a similar but less pronounced inward shift. These results indicate that Isw2-dependent remodeling reduces the region of accessible DNA adjacent to ORC under otherwise identical *in vitro* conditions.

Consistent with this architecture, deletion of *ISW2* or *CHD1*—remodelers that promote inward nucleosome positioning—increased MCM loading (Figure 2), linking NDR structure to licensing efficiency. Because helicase loading occurs on DNA immediately flanking the ORC-bound site, narrowing of the origin NDR provides a straightforward mechanistic explanation: nucleosome repositioning near the origin can limit productive helicase loading.

We next tested this prediction directly *in vivo* by mapping nucleosomes using MNase digestion in G2/M and G1 cells of WT, *isw2*Δ, *hst3Δ hst4Δ*, and the licensing-rescued *isw2*Δ *hst3*Δ *hst4*Δ mutant, allowing direct comparison of origin chromatin structure and licensing efficiency across genotypes and cell-cycle stages.

Our orientation and alignment strategy allowed nucleosomes flanking the origin to be distinguished relative to the ORC binding site. As shown in Figure 5, this arrangement revealed that the nucleosome on the left side of the origin, proximal to the ORC binding site, is more sharply positioned, consistent with constraints imposed by ORC binding. In contrast, the nucleosome on the right side occupies two closely spaced positions *in vivo*. One of these positions lies closer to the origin center and would therefore be expected to encroach more strongly on the region where MCM is loaded, potentially limiting helicase loading. This dual positioning is less apparent in *in vitro* reconstructions that isolate the action of individual remodelers together with ORC. *In vivo*, nucleosome organization likely reflects the combined effects of multiple remodelers and additional DNA-binding factors, such as transcription factors, that impose positioning constraints.

**Figure 5.**
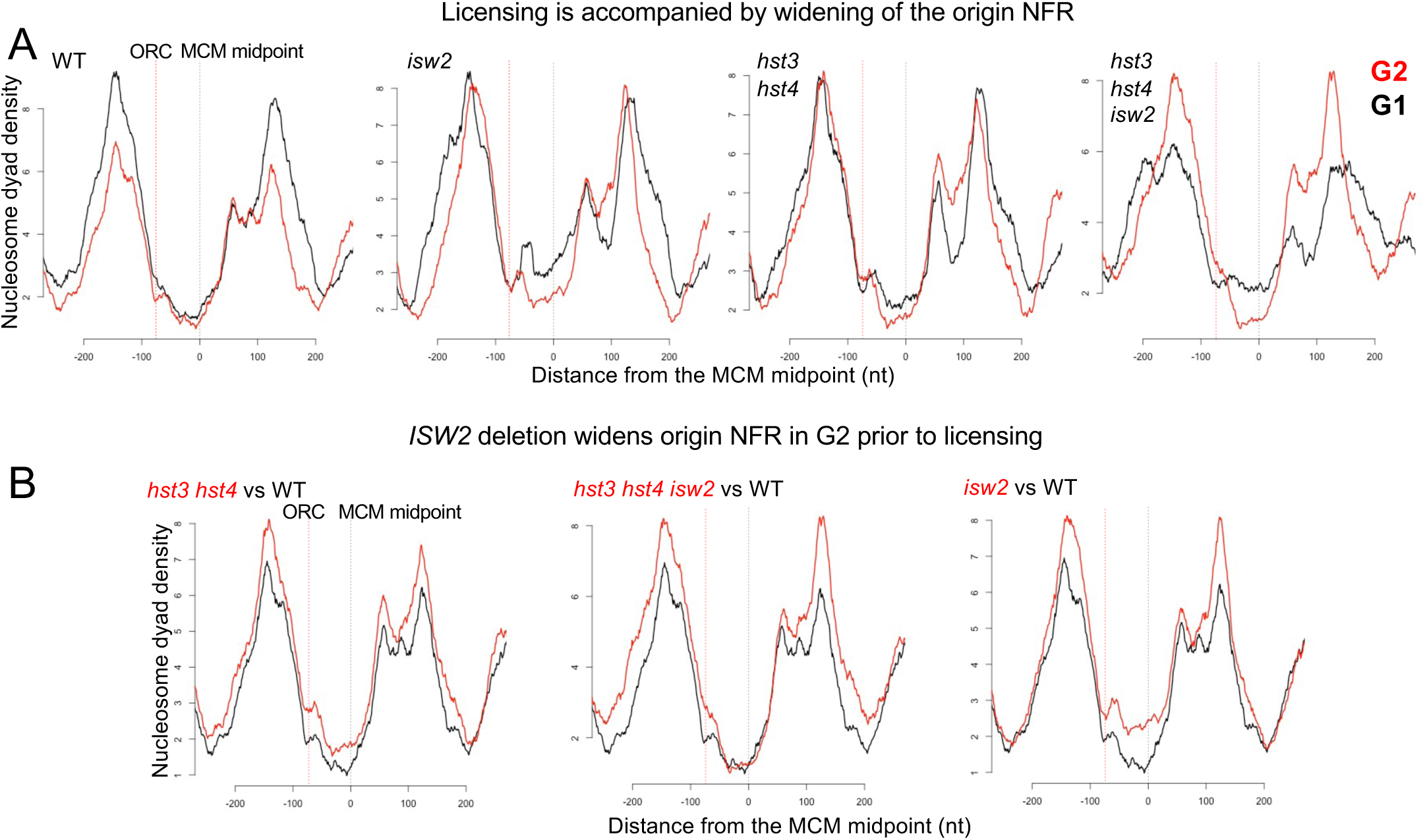
ISW2 deletion widens the origin NDR prior to licensing. (A) Nucleosome dyad density profiles derived from MNase digestion in G2/M (red) and G1 (black) cells for the indicated genotypes across 91 oriented replication origins. Profiles are aligned to the MCM midpoint and oriented as in Figures 3 and 4. Vertical dashed lines indicate the ORC binding site and the MCM midpoint. In wild-type cells, licensing in G1 is accompanied by outward repositioning of nucleosomes flanking the origin NDR, indicating expansion of the NDR during helicase loading. This repositioning is modestly enhanced in *isw2*Δ cells and reduced in *hst3*Δ *hst4*Δ mutants. In the *isw2*Δ *hst3*Δ *hst4*Δ background, nucleosome repositioning in G1 is partially restored, paralleling recovery of MCM loading. (B) Direct comparison of G2/M nucleosome organization prior to licensing. In *hst3*Δ *hst4*Δ cells, origin-flanking nucleosomes differ only subtly from wild type despite the severe licensing defect. In contrast, in the *hst3*Δ *hst4*Δ background, *ISW2* deletion produces pronounced widening of the origin NDR and outward repositioning of flanking nucleosomes in G2 prior to licensing. Because this structural change precedes helicase loading and restores licensing, these results are consistent with nucleosome encroachment limiting helicase loading.

Consistent with this origin architecture, in WT cells, licensing was accompanied by outward repositioning of nucleosomes flanking the origin NDR, indicating that helicase loading requires expansion of the origin NDR. This repositioning was modestly enhanced in isw2Δ cells and reduced in *hst3*Δ *hst4*Δ mutants and was partially restored in the isw2Δ hst3Δ hst4Δ background, paralleling recovery of licensing (Figures 5A and S8A).

Because nucleosome organization in G1 could be a consequence rather than a cause of helicase loading, we examined nucleosome positioning in G2 cells prior to licensing. In *hst3*Δ *hst4*Δ mutants, origin-flanking nucleosomes differed only subtly from wild type, showing slightly increased positional fuzziness but little change in average positioning despite the severe licensing defect. These results suggest that impaired licensing arises from subtle constraints on origin accessibility that are not fully captured by population-average nucleosome positioning.

*ISW2* deletion revealed this structural constraint most clearly in the *hst3*Δ *hst4*Δ background, where it produced pronounced widening of the origin NDR and outward repositioning of flanking nucleosomes in G2 prior to helicase loading (Figures 5B and S8B). Because this structural change precedes licensing and restores MCM loading, these results are consistent with nucleosome encroachment limiting helicase loading. Together, these findings demonstrate that Isw2-dependent remodeling narrows the origin NDR, creating a local chromatin barrier to MCM helicase loading.

Analysis of published *in vitro* datasets across the same collection of origins is consistent with this interpretation: Isw2 preferentially shifts nucleosomes inward, whereas remodelers such as Isw1 and Ino80 maintain a more expanded NDR permissive for MCM loading (Figure 4). Accordingly, both restoration of licensing in the *isw2*Δ *hst3*Δ *hst4*Δ strain and increased licensing in *isw2*Δ cells can be explained by relief of Isw2-dependent nucleosome encroachment into the origin NDR during helicase loading.

Previous biochemical work showed that H3K56 acetylation enhances nucleosome sliding by several remodelers, including Isw1 and its human orthologue Snf2h (Duan et al. 2025), raising the possibility that persistent H3K56ac directly stimulates Isw2 remodeling activity. We therefore tested whether H3K56ac alters Isw2 activity using the same *in vitro* nucleosome sliding assay, with Snf2h as a positive control. In contrast to Snf2h, which displayed increased sliding activity on H3K56ac-containing nucleosomes, Isw2 activity was not robustly stimulated by H3K56 acetylation (Figure 6). Because Isw2 promotes nucleosome encroachment into the origin NDR and its catalytic activity is required for licensing inhibition, these results suggest that persistent H3K56 acetylation does not impair licensing by strongly stimulating Isw2 remodeling activity but instead acts through changes in the chromatin context in which remodeling occurs.

**Figure 6.**
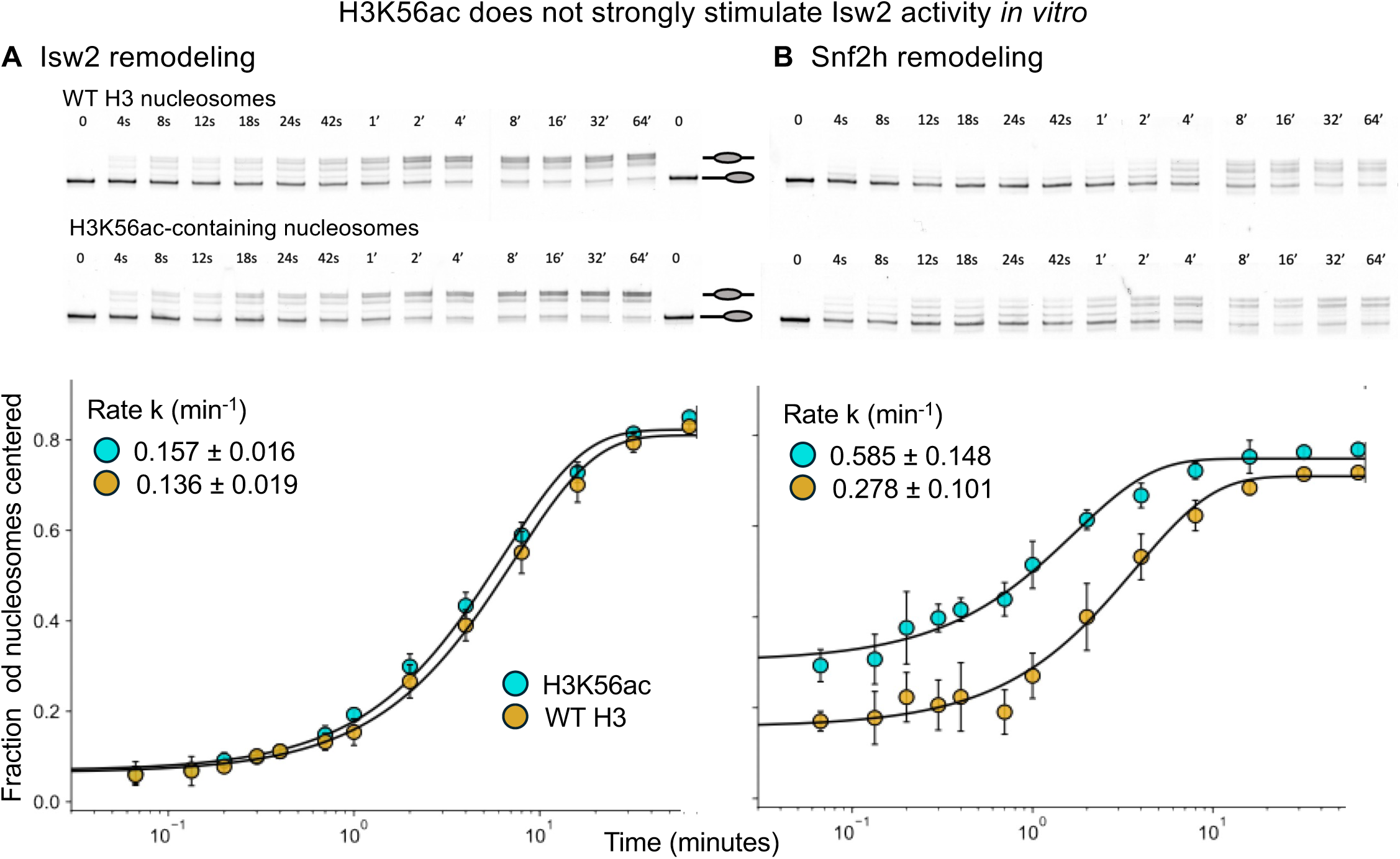
H3K56ac does not strongly stimulate Isw2 activity *in vitro*. (A) *In vitro* nucleosome sliding assays were performed using recombinant Isw2 and reconstituted nucleosomes containing either wild-type histone H3 or H3K56-acetylated H3 (H3K56ac). The fraction of centered nucleosomes was quantified over time and fit by nonlinear regression to derive apparent rate constants (k, min⁻¹). Isw2 catalytic rates were similar on wild-type and H3K56ac nucleosomes (k = 0.136 ± 0.019 min⁻¹ and 0.157 ± 0.016 min⁻¹, respectively), indicating that H3K56 acetylation does not strongly enhance Isw2 remodeling activity *in vitro*. A representative experiment is shown; rate constants were derived from two independent experiments. (B) As a positive control, the human ISWI orthologue Snf2h was assayed under identical conditions. Snf2h displayed higher sliding activity on H3K56ac nucleosomes than on wild-type nucleosomes (k = 0.585 ± 0.148 min⁻¹ versus 0.278 ± 0.101 min⁻¹), consistent with H3K56ac-dependent stimulation of remodeling activity under these assay conditions.

Together, these findings support a model in which post-replicative chromatin maturation promotes licensing of replication origins by preventing Isw2-dependent nucleosome encroachment into the origin NDR, thereby removing a chromatin-imposed barrier to MCM helicase loading.

## Discussion

Origin licensing has traditionally been viewed as a gradual process unfolding throughout G1, constrained primarily by cell-cycle control and the availability of licensing factors. Under our experimental conditions, however, most origin licensing occurred in a rapid, synchronous burst at mitotic exit, before most cells had entered G1. This timing prompted us to investigate chromatin at replication origins, revealing that post-replicative chromatin maturation during the preceding S phase establishes the capacity of origins to support helicase loading and, more broadly, that origin chromatin architecture constrains the overall extent of licensing even under otherwise permissive conditions.

A defining feature of post-replicative chromatin maturation is the transient acetylation of histone H3 on lysine 56 (H3K56ac), which is deposited on newly synthesized histones during replication-coupled nucleosome assembly by the histone acetyltransferase Rtt109 (Masumoto et al. 2005; Xu et al. 2005; Han et al. 2007). H3K56 acetylation is associated with increased nucleosome mobility and enhanced activity of ATP-dependent chromatin remodelers, creating a remodeling-prone chromatin state on newly replicated DNA (Kaplan et al. 2008; Duan et al. 2025). This state is normally resolved by Hst3- and Hst4-mediated deacetylation at the end of S phase, which stabilizes nucleosome positioning and restores mature chromatin structure (Celic et al. 2006; Maas et al. 2006; Han et al. 2007; Thaminy et al. 2007). Our results indicate that failure to properly terminate this maturation program, due to persistent H3K56 acetylation, leads to excessive remodeling of origin-proximal chromatin that inhibits MCM loading.

Consistent with this interpretation, deletion of the *ISW2* restored licensing in *hst3*Δ *hst4*Δ cells. Strikingly, loss of *ISW2* also increased MCM loading by ∼40% in wild-type cells, indicating that its inhibitory effect is not confined to chromatin maturation mutants but operates under normal physiological conditions. Together, these observations indicate that origin chromatin architecture constrains licensing capacity.

Our analysis of ORC occupancy in G2/M and G1 cells indicates that origin licensing is restricted at a step following ORC recruitment, even under conditions permissive for licensing. In G2/M cells, ORC occupancy was indistinguishable across genotypes, indicating that chromatin maturation and Isw2 do not affect ORC recruitment. In G1, however, when CDK activity is low and licensing is normally permitted, ORC occupancy diverged systematically: retention was greatest in *hst3*Δ *hst4*Δ, intermediate in wild type, and lowest in *isw2Δ*, consistent with reduced ORC association accompanying successful MCM double-hexamer assembly (Reuter et al. 2024). Substantial ORC persists at wild-type origins under permissive conditions, indicating that ORC binding does not limit MCM loading. Instead, the graded ORC retention across genotypes indicates that an Isw2-dependent chromatin state restricts helicase loading at ORC-bound origins. Thus, CDK activity determines *when* licensing can occur, whereas origin chromatin determines its extent.

Because our data indicate that origin licensing is limited at the step of helicase loading, we considered whether chromatin architecture might constrain this reaction. Biochemical and single-molecule studies have shown that loading of two oppositely oriented MCM hexamers by a single ORC complex requires ORC repositioning across the origin and engagement with DNA flanking the ACS (Ticau et al. 2015; Coster and Diffley 2017; Feng et al. 2021; Gupta et al. 2021). Consistent with this stepwise loading mechanism, ORC bound at the ACS may support initial recruitment of MCM within a relatively confined DNA region, whereas completion of double-hexamer assembly requires repositioning onto origin-flanking DNA and is therefore particularly sensitive to nucleosome positioning that limits DNA accessibility. In this framework, nucleosome encroachment into the origin NDR would be predicted to preferentially impair completion of helicase loading. Accordingly, inward positioning of origin-proximal nucleosomes would restrict the DNA available for ORC repositioning and thereby reduce overall MCM loading. Consistent with this model, deletion of *ISW2* expands the NDR and restores licensing in chromatin maturation mutants. Critically, expansion of the origin NDR upon *ISW2* deletion was already evident in G2/M cells, before helicase loading begins, indicating that nucleosome positioning is established prior to the licensing reaction and can therefore modulate its progression. Together, these observations support a model in which Isw2-dependent remodeling drives inward encroachment of origin-proximal nucleosomes, constraining the NDR and thereby limiting helicase loading. Supporting this interpretation, published *in vitro* reconstitution studies show that nucleosome arrays organized by Isw2—and, to a lesser extent, Chd1—exhibit tighter spacing and a narrower NDR than those assembled by Ino80 or Isw1 (Azmi et al. 2017; Chacin et al. 2023). The concordance between these biochemical findings and our *in vivo* analyses indicates that the licensing phenotypes arise from direct effects of Isw2 on origin-proximal nucleosome organization rather than indirect consequences of altered transcription or global chromatin changes.

Multiple ATP-dependent chromatin remodelers can organize nucleosomes flanking replication origins *in vitro*, including Isw1, Isw2, Ino80, and Chd1 (Azmi et al. 2017; Chacin et al. 2023). *In vivo*, origin architecture therefore reflects the combined—and often opposing—activities of several remodelers. Thus, these remodelers are not functionally equivalent: Isw1 and Ino80 maintain a wider NDR, whereas Isw2 promotes inward nucleosome positioning. Licensing is therefore not limited by chromatin organization itself, but by a specific remodeling activity that imposes an inhibitory nucleosome configuration. This constraint becomes particularly evident when post-replicative chromatin remains remodeling-prone, revealing Isw2 as a principal inhibitor of helicase loading.

Our findings suggest that chromatin provides a layer of replication control distinct from canonical CDK-dependent licensing regulation. Origin licensing is generally thought to be prohibited during S phase because any licensing outside G1 is assumed to be intrinsically dangerous and prone to drive re-replication. However, this view conflates *when* licensing occurs with *where* it occurs. In principle, licensing on unreplicated DNA during S phase is not inherently problematic, because it does not create an opportunity for a second round of replication at that locus. By contrast, the pathological event is licensing on already replicated DNA, which permits aberrant re-initiation and genomic instability. Canonical mechanisms that prevent re-replication—such as ORC phosphorylation, MCM nuclear exclusion, and Cdc6 destabilization—operate at the level of the cell cycle and act uniformly across the genome, without regard to the replication status of individual loci. Our results identify a chromatin-based licensing barrier that specifically suppresses helicase loading on newly replicated DNA. In this model, H3K56 acetylation on newly replicated chromatin promotes Isw2-dependent nucleosome remodeling that inhibits MCM loading, whereas removal of H3K56ac at S-phase exit erases this inhibitory state and restores origin competence for the next round of licensing.

Together, our results show that origin licensing is governed not only by cell-cycle regulation but also by the structural state of origin chromatin (Figure 7). Even under otherwise permissive conditions, origin-proximal nucleosomes constrain helicase loading, and post-replicative chromatin maturation relieves—but does not eliminate—this constraint, thereby modulating the overall extent of origin licensing. We therefore propose that licensing is controlled by two integrated systems: CDK-dependent regulation that determines when licensing can occur, and origin chromatin architecture that determines how extensively individual origins are licensed. In this view, replication origins exist along a continuum of licensing competence defined by their local chromatin environment. Taken together, our findings establish origin chromatin architecture as a key determinant of licensing capacity and suggest that chromatin-encoded control of origin usage represents a general organizing principle of eukaryotic genome duplication.

**Figure 7.**
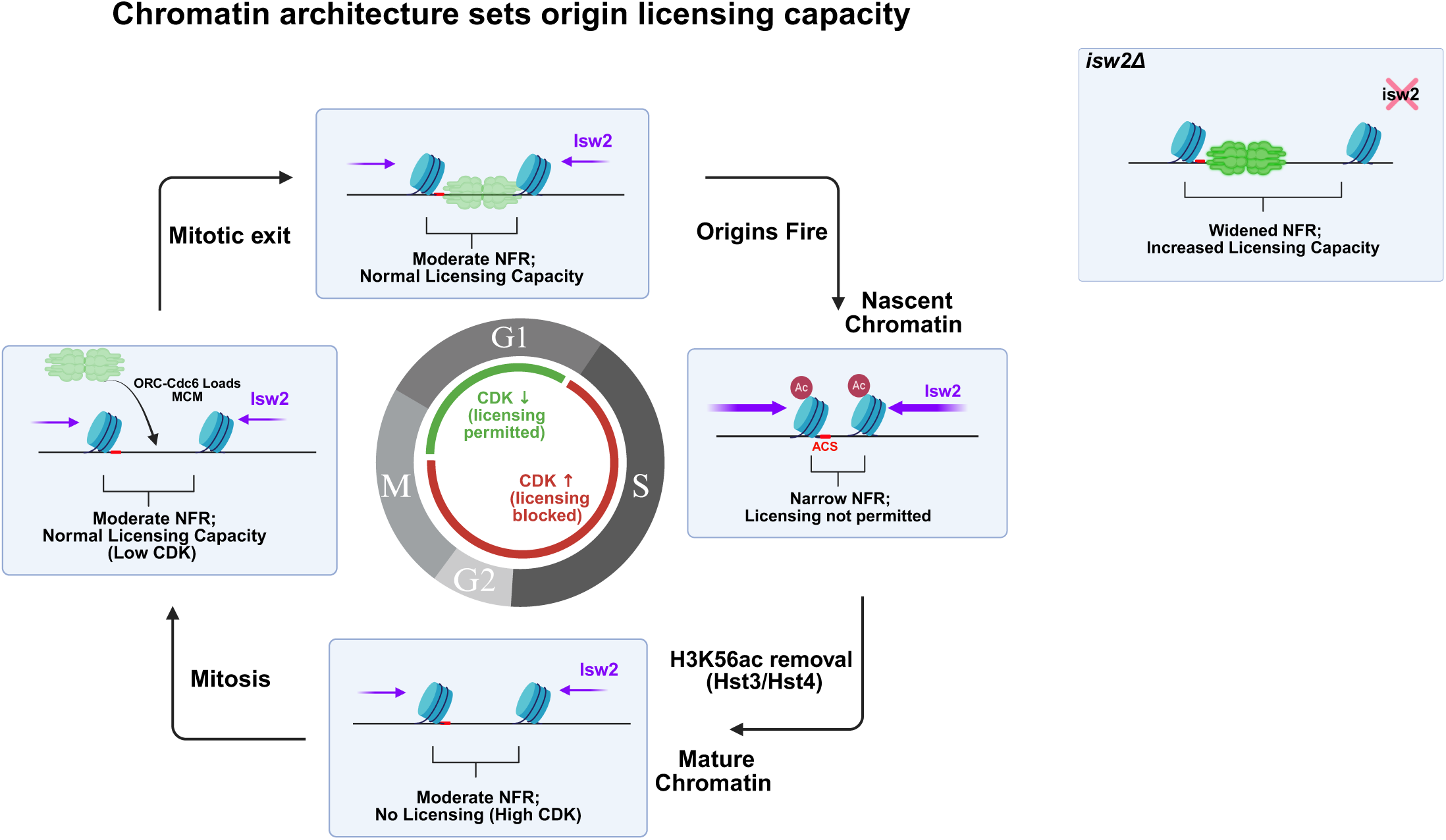
Chromatin architecture sets origin licensing capacity. Model illustrating how origin chromatin architecture and CDK activity jointly regulate replication origin licensing. During S phase, newly replicated chromatin is assembled with H3K56 acetylation and adopts a nucleosome configuration in which the origin-proximal nucleosome encroaches on the nucleosome-free region (NFR), limiting access to the helicase loading site. As chromatin matures following S phase, H3K56 acetylation is removed by Hst3 and Hst4, allowing nucleosomes flanking the origin to reposition and partially restore the NFR. At mitotic exit, CDK activity declines, permitting helicase loading. Under these licensing-permissive conditions, the width of the origin NFR determines how efficiently MCM double hexamers are loaded. Isw2 acts to reposition origin-flanking nucleosomes in a manner that limits NFR width and thereby constrains licensing capacity. In cells lacking Isw2, the NFR expands, allowing increased MCM loading. Thus, origin licensing is controlled by two integrated systems: CDK-dependent regulation that determines when licensing can occur and chromatin architecture that determines how extensively origins are licensed.

## Methods

### Yeast strains and growth conditions

*S. cerevisiae* strains used in this study are listed in Table S1, plasmids in Table S2, and primers in Table S3. All strains are in the S288C genetic background. Gene deletions and epitope tagging were generated using standard PCR-based homologous recombination. The *isw2*-K215R allele was generated using a pop-in/pop-out gene replacement strategy with an integrating plasmid (Gelbart et al., 2005). Cells were grown at 30 °C in YPD medium (1% yeast extract, 2% peptone, 2% glucose) unless otherwise indicated.

### Cell-cycle synchronization

For cell-cycle time-course experiments, cultures were grown to mid-log phase and arrested in G2/M by addition of nocodazole (15 µg mL⁻¹) for 3 h. Arrest efficiency was verified by microscopy and DNA-content analysis. Cells were released from arrest by washing twice with fresh YPD medium and resuspending in nocodazole-free medium supplemented with α-factor (5 µg mL⁻¹). Samples were collected at the indicated time points for ChEC assays, immunoblotting, and flow cytometry.

### Chromatin endogenous cleavage (ChEC–seq)

ChEC–seq was performed as previously described (Foss et al. 2019; Foss et al. 2021). *S. pombe* cells expressing an Mcm2–MNase fusion protein in log phase were added as spike-in controls prior to ChEC. Cells expressing MNase fusion proteins were permeabilized with digitonin and MNase cleavage was induced by addition of CaCl₂. Reactions were carried out for the indicated times and terminated by addition of SDS and EDTA-containing stop buffer. Undigested control samples were collected prior to calcium addition.

Following cleavage, samples were treated with proteinase K and DNA was purified by phenol–chloroform extraction and ethanol precipitation. MNase digestion patterns were assessed by agarose gel electrophoresis prior to library preparation. Sequencing libraries were prepared from total DNA without size selection as previously described (Foss et al. 2019; Foss et al. 2021).

Reads mapping to the *S. pombe* genome were used to calculate normalization factors between samples. *S. cerevisiae* read counts were scaled according to the relative abundance of *S. pombe* reads to correct for variation in immunoprecipitation efficiency and sequencing depth. Spike-in normalization was not applied to ORC chromatin immunoprecipitation experiments.

### Sequence alignment and spike-in normalization

Sequencing reads were aligned using BWA to both the *Saccharomyces cerevisiae* (sacCer3) and *S. pombe* (ASM294v2) genomes (Li and Durbin 2009). Reads mapping to 1966 1 kb segments in the *S. pombe* genome corresponding to previously mapped MCM loading sites were used to calculate normalization factors between samples, and *S. cerevisiae* read counts were scaled according to the relative abundance of *S. pombe* spike-in reads to correct for variation in MNase cleavage efficiency, DNA recovery, and sequencing depth.

The goal of this normalization was to express *S. cerevisiae* MCM loading, which varied across the time course, relative to the invariant MCM loading observed in log-phase *S. pombe* cells.

### Quantification of MCM loading kinetics

ChEC–seq signal reflects cleavage events generated by MNase fused to MCM proteins at sites of helicase loading. Aggregate profiles and heatmaps were generated by aligning sequencing reads relative to replication origin coordinates. For genome-wide analyses, approximately 326 origins were aligned relative to the autonomously replicating sequence (ACS). For analyses of oriented origins, a curated set of 91 origins with defined ORC orientation was aligned relative to the midpoint of the MCM signal distribution.

To quantify licensing kinetics, MCM signal at each origin was measured across the time course and normalized to the maximal signal observed for that origin. The time required to reach half-maximal loading (t½) was calculated for each origin and used to order origins for heatmap visualization.

### ChEC–qPCR

Origin licensing at individual loci was quantified using ChEC–qPCR as described and validated previously (Foss et al. 2021). Following MNase activation as described above, DNA was purified and analyzed by quantitative PCR using primers spanning replication origins. MNase cleavage at licensed origins disrupts the PCR amplicon, reducing qPCR signal in proportion to the fraction of origins that have loaded an MCM double hexamer. Signals were normalized to a non-origin control locus and expressed relative to uncleaved control samples.

### Chromatin immunoprecipitation (Ch IP)

Chromatin immunoprecipitation was performed as previously described (Lichauco et al. 2025), with some modifications. For spike-in normalization in Mcm2 immunoprecipitation experiments, *Schizosaccharomyces pombe* cells expressing FLAG-tagged Mcm2 were mixed with *Saccharomyces cerevisiae* cells expressing FLAG-tagged proteins prior to crosslinking. Cells were crosslinked with 2% formaldehyde for 30 minutes and quenched with glycine.

Chromatin was prepared using two approaches. In the first, cells were lysed by glass bead disruption and chromatin was fragmented by sonication to generate DNA fragments of approximately 200–300 bp. In the second approach, nuclei were prepared from spheroplasted cells and chromatin was fragmented by digestion with micrococcal nuclease (MNase). An aliquot of fragmented chromatin was retained as an input control.

Immunoprecipitations were performed using Dynabeads Protein G pre-bound with anti-FLAG antibody (Sigma). After washing, bound chromatin was eluted and crosslinks were reversed. DNA was purified using a MinElute PCR purification kit and used for Illumina library preparation and sequencing.

For analysis of ORC occupancy, sequencing reads were analyzed using a curated set of 91 replication origins with defined ORC orientation aligned relative to the midpoint of the MCM signal distribution determined from ChEC experiments.

For Mcm2 ChIP experiments, reads mapping to the *S. pombe* genome were used to calculate normalization factors between samples. *S. cerevisiae* read counts were scaled according to the relative abundance of *S. pombe* reads to correct for variation in immunoprecipitation efficiency and sequencing depth. Spike-in normalization was not applied to ORC chromatin immunoprecipitation experiments.

### MNase mapping of nucleosome positioning

MNase-seq was performed as previously described (Foss et al. 2019). Briefly, log-phase cells were synchronized with α-factor, crosslinked with formaldehyde, converted to spheroplasts, and nuclei were digested with MNase. MNase digestion conditions were selected to yield the expected mono-, di-, and tri-nucleosome protected DNA fragments, as assessed by gel electrophoresis prior to library preparation. DNA was purified after reversal of crosslinks and used for sequencing library preparation.

### Immunoblotting

Protein extracts were prepared using the alkaline lysis method (Kushnirov, 2000). Cells were treated with NaOH to permeabilize the cell wall, resuspended in SDS sample buffer containing β-mercaptoethanol, and boiled prior to SDS–PAGE. Proteins were separated by SDS–PAGE and transferred to nitrocellulose membranes. Membranes were probed with antibodies against Clb2, Clb5, and Sic1, followed by horseradish peroxidase–conjugated secondary antibodies. Signals were detected using chemiluminescence.

### Flow cytometry

Cell-cycle progression was monitored by DNA-content analysis. Cells were fixed in 70% ethanol, treated with RNase A and proteinase K, and stained with propidium iodide. DNA content was measured using a flow cytometer and analyzed using FlowJo.

### Nucleosome sliding assays

Recombinant *S. cerevisiae* Isw2 and human SNF2h were purified as described in Duan et al. 2025. Nucleosomes were reconstituted on the Widom 601 positioning sequence using recombinant *Xenopus laevis* histones; H3K56-acetylated histone H3 was obtained from the Histone Source (Colorado State University). Sliding reactions contained 40 nM nucleosomes and 200 nM remodeler and were initiated by addition of ATP. Reactions were quenched at the indicated time points and resolved by native PAGE, and nucleosome repositioning was quantified as the fraction of centered nucleosomes over time as described previously (Duan et al. 2025). Apparent rate constants were obtained by nonlinear regression.

### Quantification and Statistical Analysis

Statistical analyses were performed using paired two-tailed t-tests unless otherwise indicated. Differences were considered statistically significant when p < 0.05. Data are presented as mean ± SEM. Statistical details for individual experiments are provided in the corresponding figure legends.

## Data availability

Sequencing data generated in this study have been deposited in the NCBI Sequence Read Archive (SRA) under BioProject accession PRJNA1452552.

## Code availability

Custom scripts used for analysis are available from the corresponding author upon request.

## Author contributions

E.J.F. and A.B. conceived the study. E.J.F. performed the computational analyses. A.G., A.U., B.L., G.D.B., H.F., I.M.N., S.M., T.T. and Z.Z. contributed experimental data, reagents, methodology, and conceptual input and analysis. E.J.F. and A.B. wrote the manuscript with input from all authors.

## Competing interests

The authors declare no competing interests.

## Supporting information

Foss_et_al_Supplemental_Tables_S4_to_S11

## Acknowledgements

We thank members of the Bedalov laboratory for helpful discussions. This work was supported by the National Institutes of Health (R01GM117446 to A.B.; R35GM139429 to T.T.; R01GM084192 to G.D.B.). Support for this work was provided in part by the David and Deborah Lycette Endowed Chair for Cancer Research (to T.T.)

## Supplemental Figures

**Figure S1.**
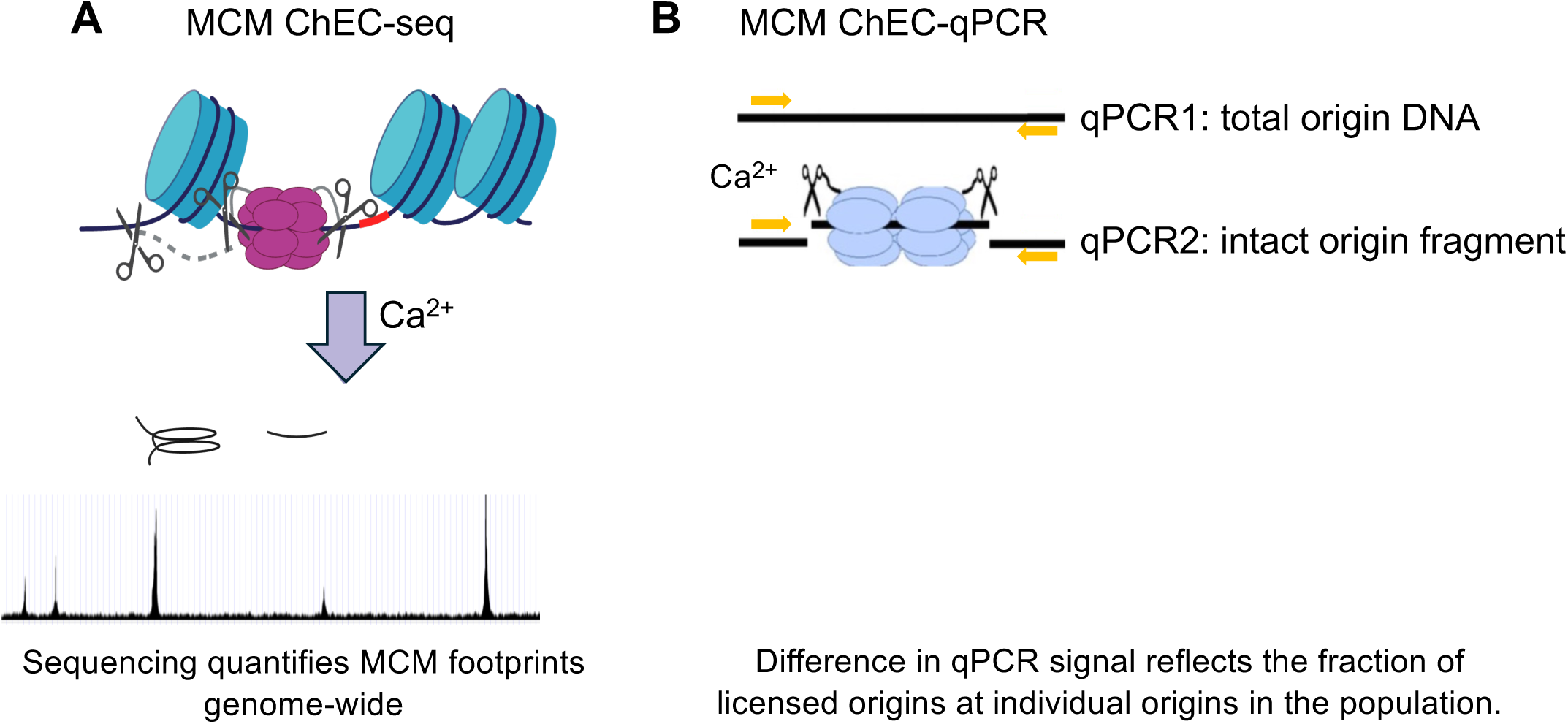
Principle of the MCM–ChEC assays used to measure origin licensing. (A) MCM–ChEC-seq. Micrococcal nuclease (MNase) fused to an MCM subunit cleaves DNA adjacent to loaded MCM double hexamers upon addition of Ca²⁺. Sequencing of the resulting fragments quantifies MCM footprints genome-wide, providing a measure of helicase loading at replication origins. Libraries were prepared without size selection, ensuring retention of small MNase fragments; larger fragments are disfavored during library preparation due to their size. (B) MCM–ChEC-qPCR. MNase cleavage at licensed origins disrupts the PCR amplicon spanning the cleavage site. Quantitative PCR using origin-specific primers amplifies only intact origin DNA; cleavage therefore reduces the qPCR signal in proportion to the fraction of origins that have undergone MNase cleavage. Because MNase cleavage occurs only at origins that have loaded an MCM double hexamer, the decrease in qPCR signal provides a quantitative measure of the fraction of licensed origins in the population.

**Figure S2.**
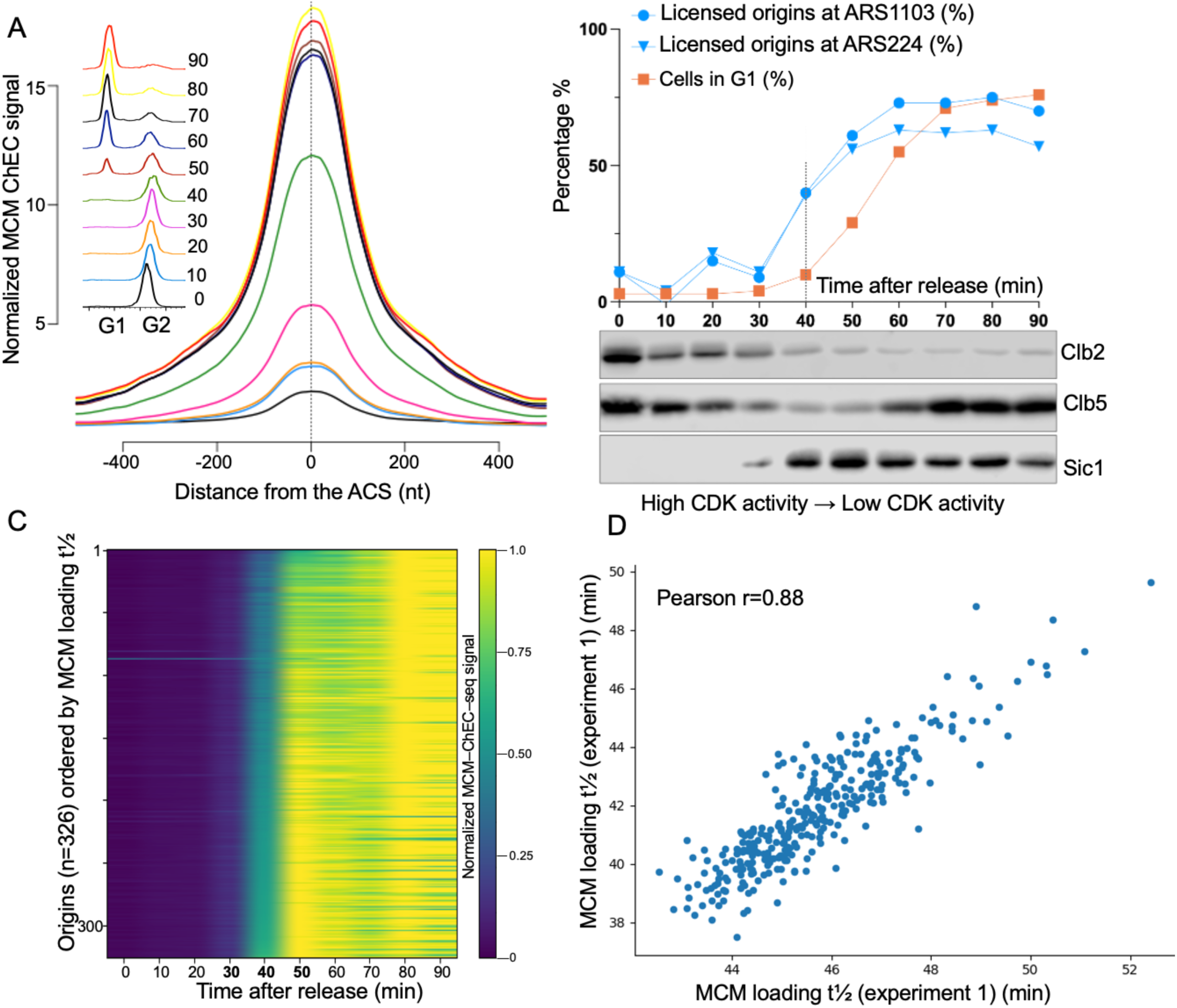
Independent replicate of the replication origin licensing time course. (A) Aggregate MCM–ChEC–seq signal across origins following release from nocodazole arrest. Normalized MCM–ChEC signal averaged across 326 annotated replication origins and aligned relative to the ACS is shown for the indicated times after release. As observed in the experiment shown in Figure 1, MCM signal is low in G2/M and increases sharply within ∼20–40 min after release, reaching near-maximal levels by ∼50–60 min. Insets show representative DNA-content profiles corresponding to each time point. (B) Quantification of licensing and CDK regulatory markers during the same time course. The fraction of licensed origins at two origins (ARS1103 and ARS224) was measured by ChEC–qPCR (blue) and compared with the fraction of cells in G1 determined by flow cytometry (orange). Immunoblot analysis of Clb2 and Clb5 together with the CDK inhibitor Sic1 shows rapid loss of B-type cyclins and accumulation of Sic1 coincident with the onset of MCM loading, consistent with licensing occurring as CDK activity declines. qPCR results are available in Supplemental Table S5. (C) Genome-wide kinetics of helicase loading. Heatmap of normalized MCM–ChEC–seq signal across 326 origins, aligned relative to the ACS as in (A) and ordered by the time to half-maximal MCM loading (t½). As in Figure 1, most origins acquire MCM signal within the same narrow time window, demonstrating highly synchronous genome-wide helicase loading. MCM-ChEC signal and t½ values for individual origins are provided in Supplemental Table S4. (D) Reproducibility of origin-specific licensing kinetics. Comparison of the time to half-maximal MCM loading (t½) for each origin between two independent experiments. Each point represents one origin (n = 326). The strong correlation (Pearson r = 0.88) indicates that origin-specific licensing kinetics are highly reproducible despite a modest offset in absolute timing between experiments.

**Figure S3.**
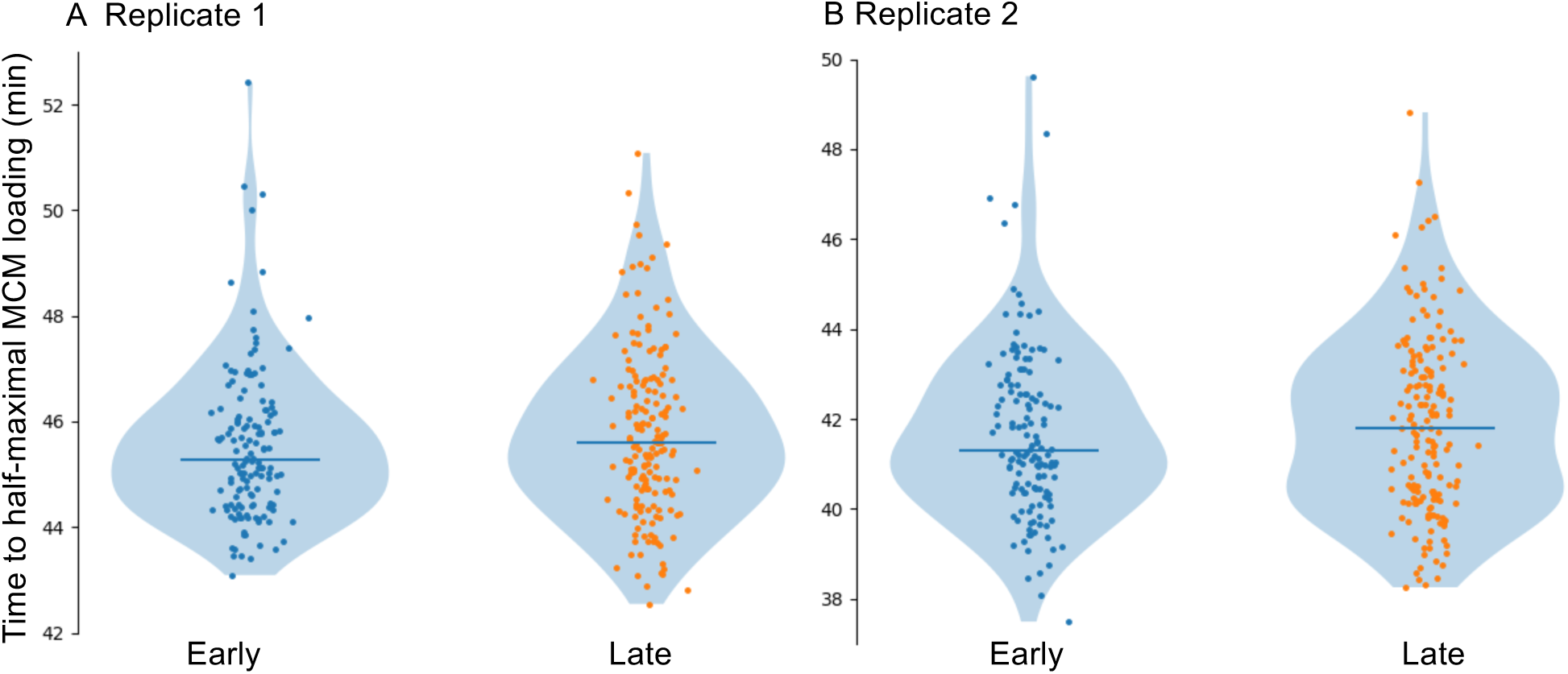
Early- and late-replicating origins exhibit similar licensing kinetics. (A) Distribution of the time to half-maximal MCM loading (t½) for origins previously classified as early-or late-replicating in the experiment shown in Figure 1. Each point represents one origin, and violin plots show the distribution of t½ values within each class. Horizontal lines indicate the mean. Classification of early and late origins and t½ values for individual origins are provided in Supplemental Table S4. (B) Same analysis as in (A) for the independent replicate experiment shown in Figure S2. Early- and late-replicating origins exhibit largely overlapping distributions of licensing kinetics in both experiments, indicating that replication timing classes do not differ substantially in the timing of MCM loading.

**Figure S4.**
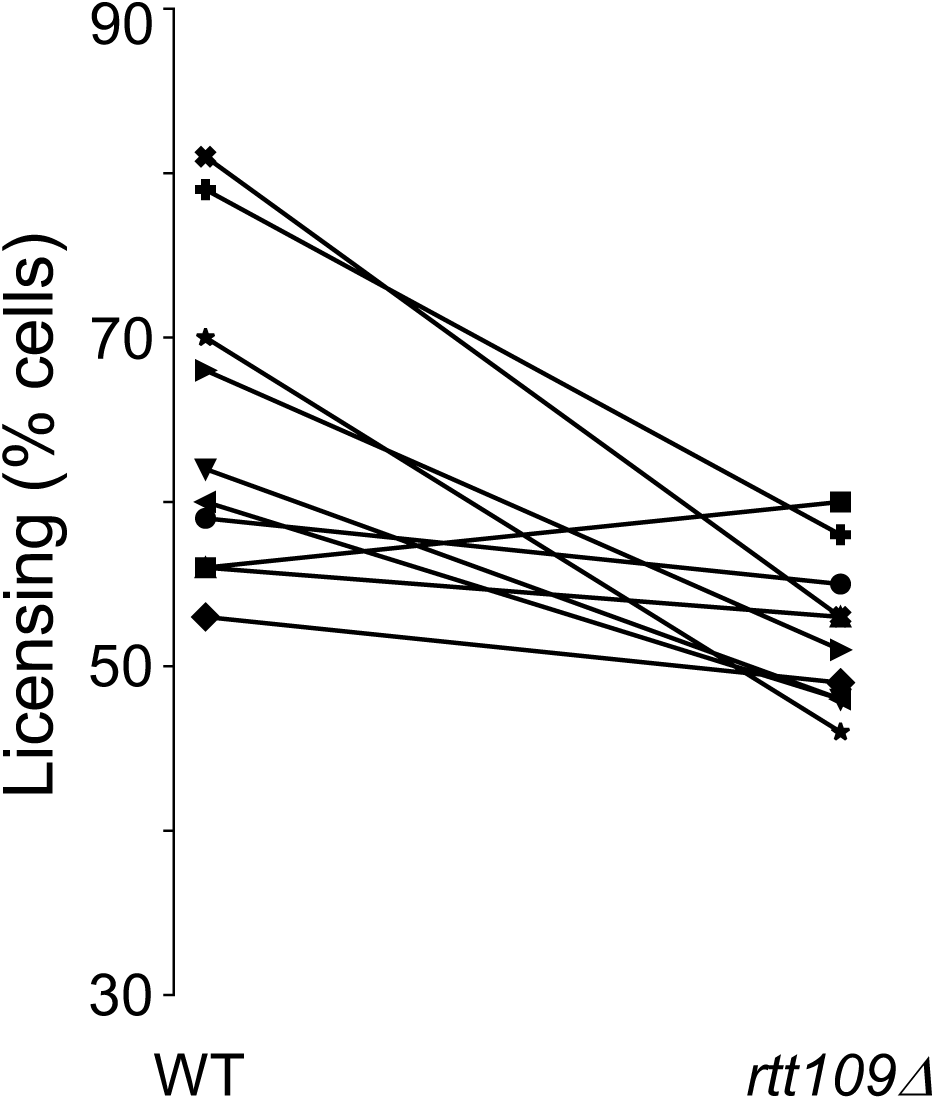
Independent replicate confirming reduced origin licensing in *rtt109*Δ cells. MCM loading was measured by ChEC–qPCR at ten replication origins in wild-type and *rtt109*Δ cells in an independent experiment. As observed in Figure 2A, loss of H3K56 acetylation in *rtt109*Δ cells caused a modest but reproducible reduction in licensing relative to wild type (mean −19%; range −35% to +7%; paired two-tailed t-test p = 0.0046). Each line represents an individual origin. Individual qPCR values are provided in Supplemental Table S8.

**Figure S5.**
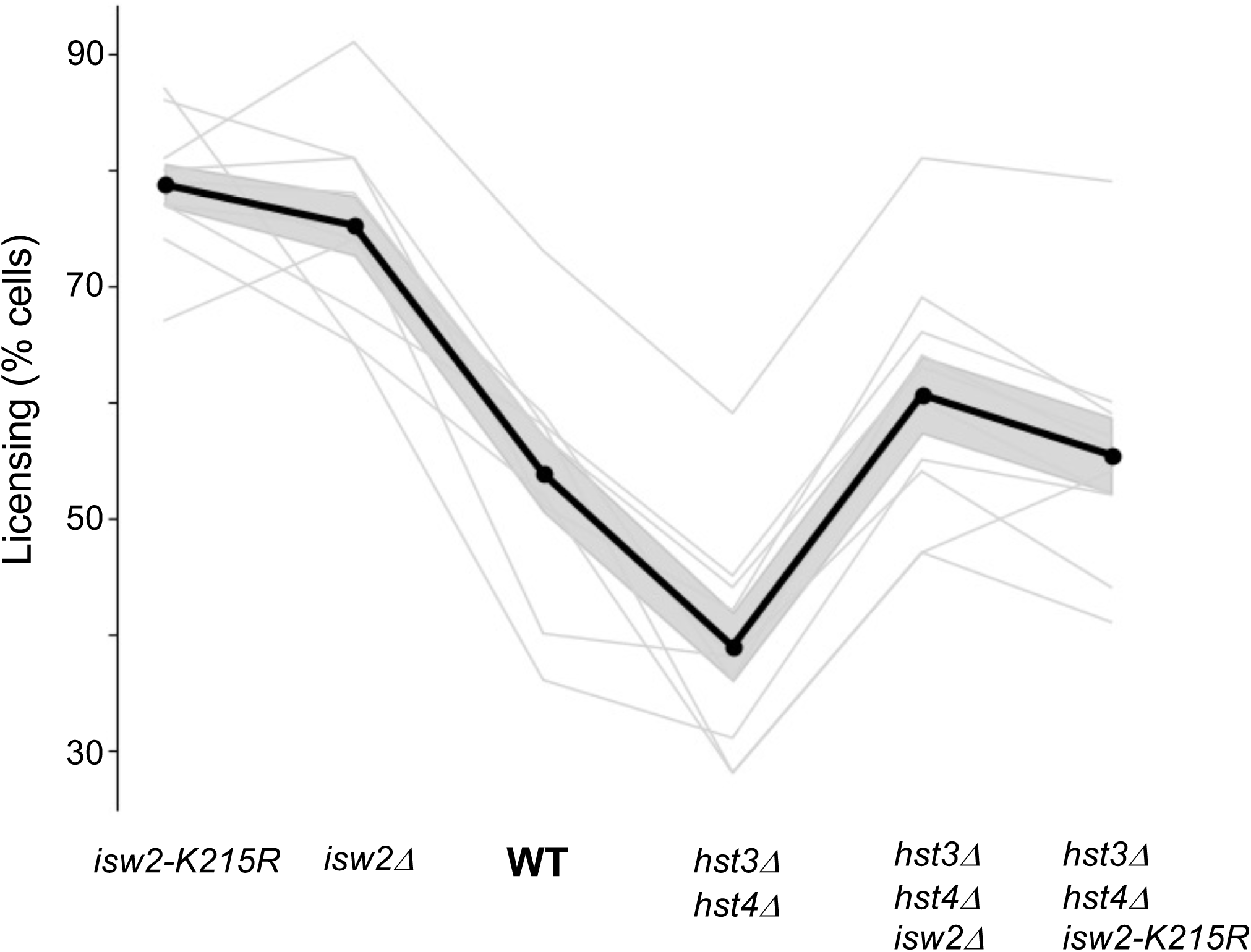
Independent biological replicate of the experiment shown in. **Figure 2D**. MCM loading was measured by ChEC–qPCR at ten replication origins in the indicated strains in an independent biological replicate. The same pattern observed in Figure 2 was reproduced: deletion of *ISW2* increased licensing relative to wild type, whereas *hst3*Δ *hst4*Δ cells exhibited a pronounced licensing defect that was strongly suppressed by deletion of *ISW2*. The ATPase-dead *isw2*-K215R allele phenocopied *isw2*Δ in both wild-type and *hst3*Δ *hst4*Δ backgrounds, indicating that catalytic remodeling activity of Isw2 is required for inhibition of origin licensing. Thin gray lines represent individual origins; the thick black line indicates the mean across origins, and the shaded region denotes ± SEM. Individual qPCR values are provided in Supplemental Table S6.

**Figure S6.**
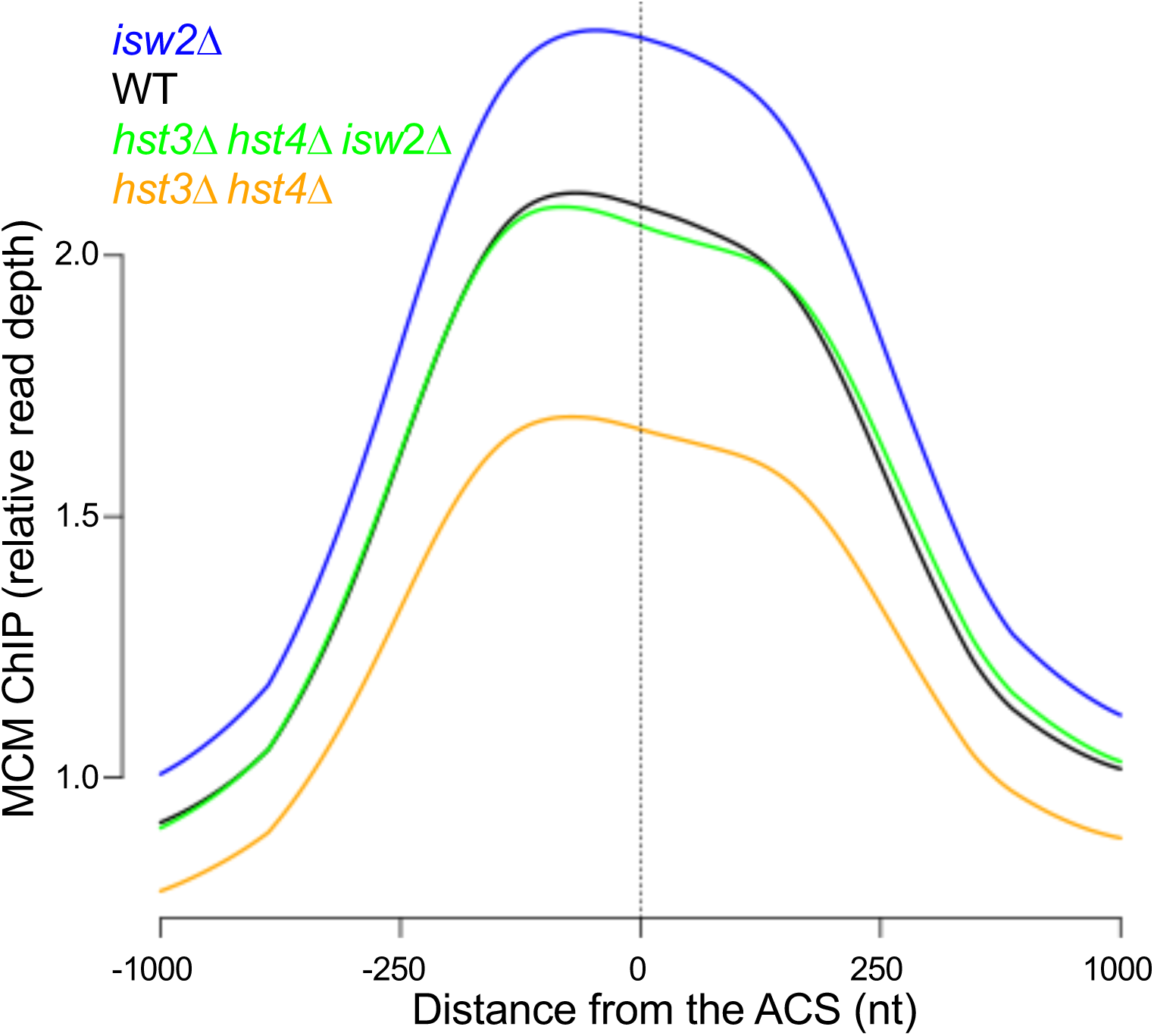
Genome-wide MCM ChIP confirms licensing differences across genotypes. Average MCM ChIP signal across 326 replication origins aligned relative to the ARS consensus sequence (ACS) in the indicated genotypes (WT, *isw2*Δ, *hst3*Δ *hst4*Δ, and *isw2*Δ *hst3*Δ *hst4*Δ). Origin-associated MCM signal is reduced in *hst3*Δ *hst4*Δ, increased in *isw2*Δ, and restored in *isw2*Δ *hst3*Δ *hst4*Δ, consistent with the licensing differences measured by MCM–ChEC. An independent biological replicate yielded highly similar results; the corresponding values for both replicates are provided in Supplemental Table S11.

**Figure S7.**
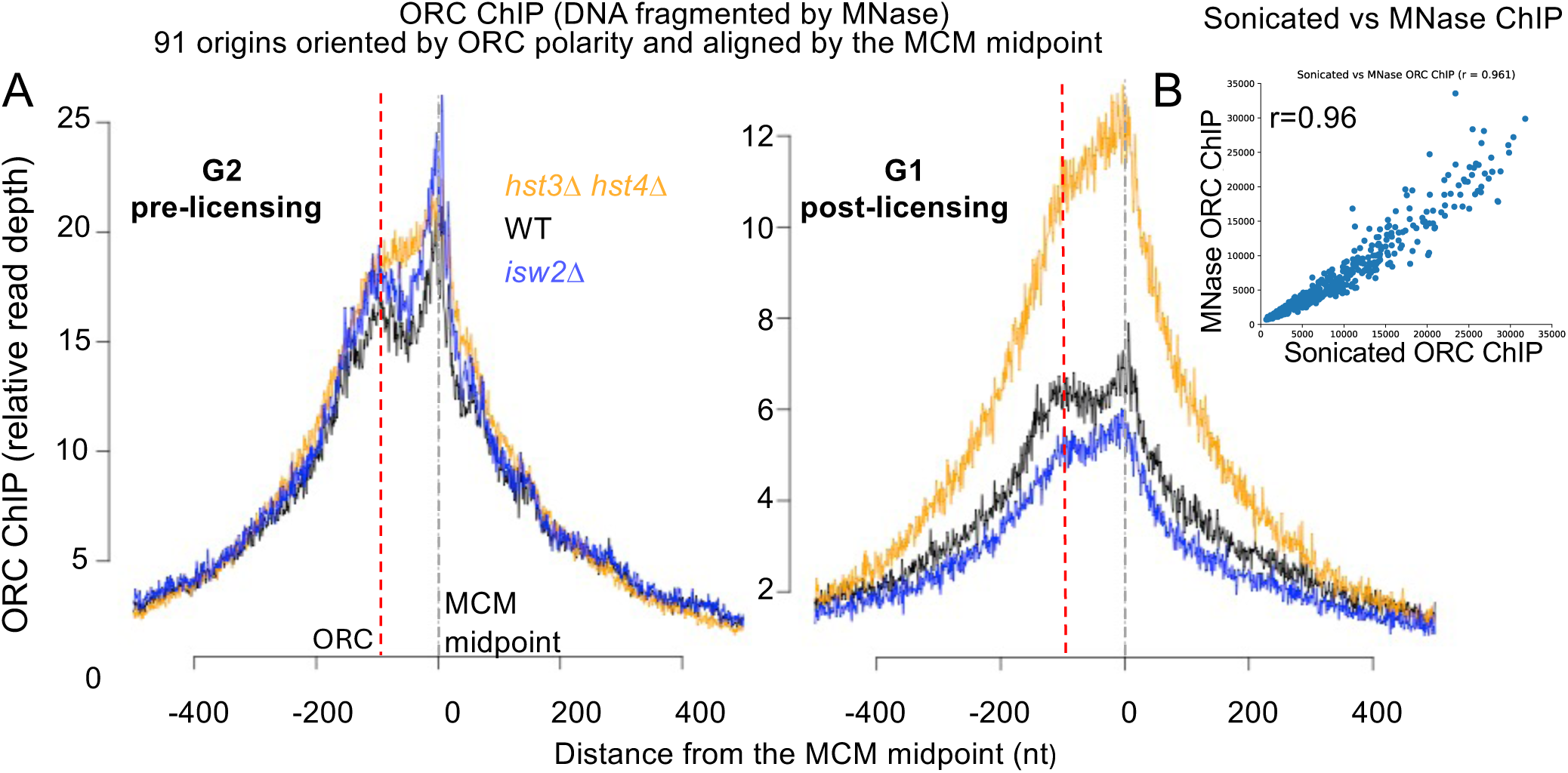
ORC ChIP profiles using MNase-fragmented chromatin. (A) Average ORC ChIP signal across 91 replication origins oriented by ORC polarity and aligned by the MCM loading midpoint in G2/M (pre-licensing) and G1 (post-licensing) cells of the indicated genotypes (WT, *isw2*Δ, and *hst3*Δ *hst4*Δ). Chromatin was fragmented using exogenous MNase digestion. As in the sonicated ChIP experiment shown in Figure 3, the ORC peak lies to the left of the MCM midpoint, consistent with the orientation of the curated origin set. Although the contours of the profiles differ from those obtained with sonicated chromatin, reflecting differences in fragmentation, the relative ORC occupancy patterns across genotypes are preserved. Relative ORC ChIP signal at each of the 91 origins for both fragmentation methods is provided in Supplemental Table S10. (B) Correlation between ORC ChIP signals obtained from sonicated and MNase-fragmented chromatin across all origins and conditions (Pearson *r* = 0.96, *N* = 546), demonstrating strong agreement between the two fragmentation methods.

**Figure S8.**
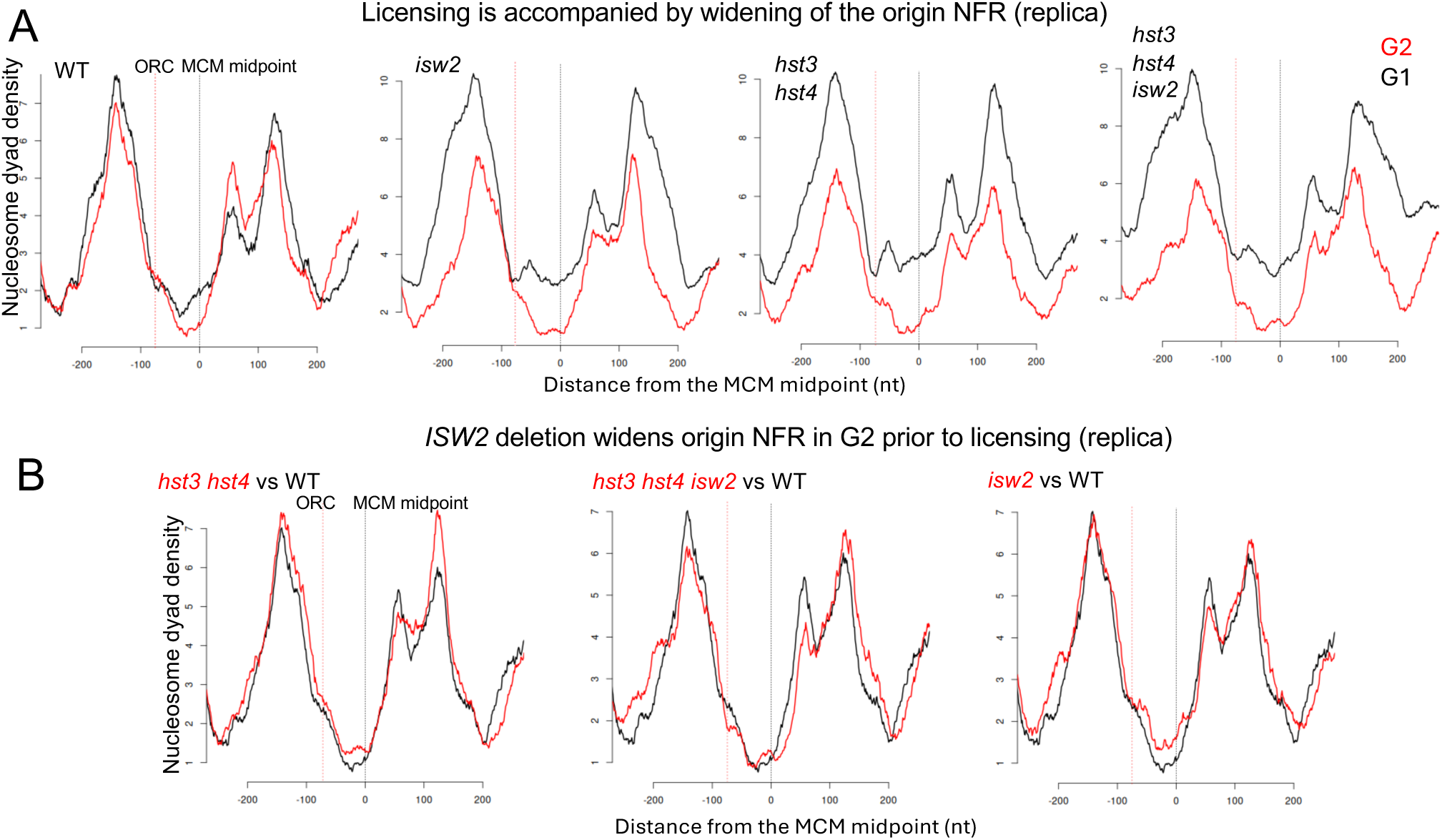
Independent biological replicate of nucleosome mapping at replication origins. (A) Nucleosome dyad density profiles derived from MNase digestion in G2/M (red) and G1 (black) cells for the indicated genotypes across 91 oriented replication origins. Profiles are aligned to the MCM midpoint and oriented as in Figures 3–5. Vertical dashed lines indicate the ORC binding site and the MCM midpoint. As in Figure 5A, licensing in wild-type cells is accompanied by outward repositioning of nucleosomes flanking the origin NDR during G1. (B) Direct comparison of nucleosome organization in G2/M prior to licensing. As in Figure 5B, deletion of *ISW2* widens the origin NDR and repositions flanking nucleosomes outward in the *hst3*Δ *hst4*Δ background. These results reproduce the nucleosome architecture changes observed in Figure 5.

## Supplemental Tables

**Supplemental Table S1.** Yeast strains used in this study.

| Strain ID | Genotype | Source |
| --- | --- | --- |
| 16652 | MATa his3 leu2 ura3 met15 | This study |
| 16964 | MATa his3 leu2 ura3 met15 hmlα $\Delta$::NAT MCM2-3xFLAG-MNase::KanMX | This study |
| 16991 | GP2 h<sup>-</sup> (L972) wild-type *Schizosaccharomyces pombe* | Gift from G. Smith |
| 17000 | GP2 h<sup>-</sup> (L972) MCM2-3xFLAG-MNase::KanMX | This study |
| 17862 | MATa his3 leu2 ura3 met15 hmlα $\Delta$::NAT MCM2-3xFLAG-MNase::KanMX isw2Δ::Hyg | This study |
| 17868 | MATa his3 leu2 ura3 met15 hmlα $\Delta$::NAT MCM2-3xFLAG-MNase::KanMX isw1Δ::Hyg | This study |
| 17870 | MATa his3 leu2 ura3 met15 hmlα $\Delta$::NAT MCM2-3xFLAG-MNase::KanMX chd1Δ::Hyg | This study |
| 17876 | MATa his3 leu2 ura3 met15 hmlα $\Delta$::NAT MCM2-3xFLAG-MNase::KanMX nhp10Δ::Hyg | This study |
| 17878 | MATa his3 leu2 ura3 met15 hmlα $\Delta$::NAT MCM2-3xFLAG-MNase::KanMX rtt109Δ::Hyg | This study |
| 17887 | MATa his3 leu2 ura3 met15 hmlα $\Delta$::NAT MCM2-3xFLAG-MNase::KanMX hst4Δ::Hyg hst3Δ::LEU2 | This study |
| 17913 | MATa his3 leu2 ura3 met15 isw2Δ::Hyg | This study |
| 17923 | MATa his3 leu2 ura3 met15 hmlα $\Delta$::NAT MCM2-3xFLAG-MNase::KanMX hst4Δ::Hyg hst3Δ::LEU2 isw2Δ::URA3 | This study |
| 17941 | MATa his3 leu2 ura3 met15 hst4Δ::Hyg hst3Δ::LEU2 isw2Δ::URA3 | This study |
| 17945 | MATa his3 leu2 ura3 met15 hst4Δ::KanMX hst3Δ::Hyg | This study |
| 17947 | MATa his3 leu2 ura3 met15 SIC1-TAP::HIS3MX6 | Open Biosystems |
| 17948 | MATa his3 leu2 ura3 met15 CLB2-TAP::HIS3MX6 | Open Biosystems |
| 17949 | MATa his3 leu2 ura3 met15 CLB5-TAP::HIS3MX6 | Open Biosystems |
| 17967 | MATa his3 leu2 ura3 met15 ORC2-6xGly-3xFLAG::KanMX | This study |
| 17976 | MATa his3 leu2 ura3 met15 ORC2-6xGly-3xFLAG::KanMX isw2Δ::HIS3 | This study |
| 17980 | MATa his3 leu2 ura3 met15 ORC2-6xGly-3xFLAG::KanMX hst3Δ::Hyg hst4Δ::NAT | This study |
| 18100 | MATa his3 leu2 ura3 met15 hmlα $\Delta$::NAT MCM2-3xFLAG-MNase::KanMX hst4Δ::Hyg hst3Δ::LEU2 isw2-K215R | This study |
| 18103 | MATa his3 leu2 ura3 met15 hmlα $\Delta$::NAT MCM2-3xFLAG-MNase::KanMX isw2-K215R | This study |

**Supplemental Table S2.** Plasmids used in this study.

| **Plasmid ID** | **Description** | **Source** |
| --- | --- | --- |
| pGZ108 | pFA6a-3FLAG-MNase-kanMX6, Addgene #70231 | Zentner et al., 2015 |
| pGZ109 | pFA6a-3FLAG-MNase-HIS3MX6, Addgene #70232 | Zentner et al., 2015 |
| pRS406-ISW2-GRT | K215R mutation in the ATPase domain of ISW2 | Gelbart et al., 2005 |

**Supplemental Table S3.**
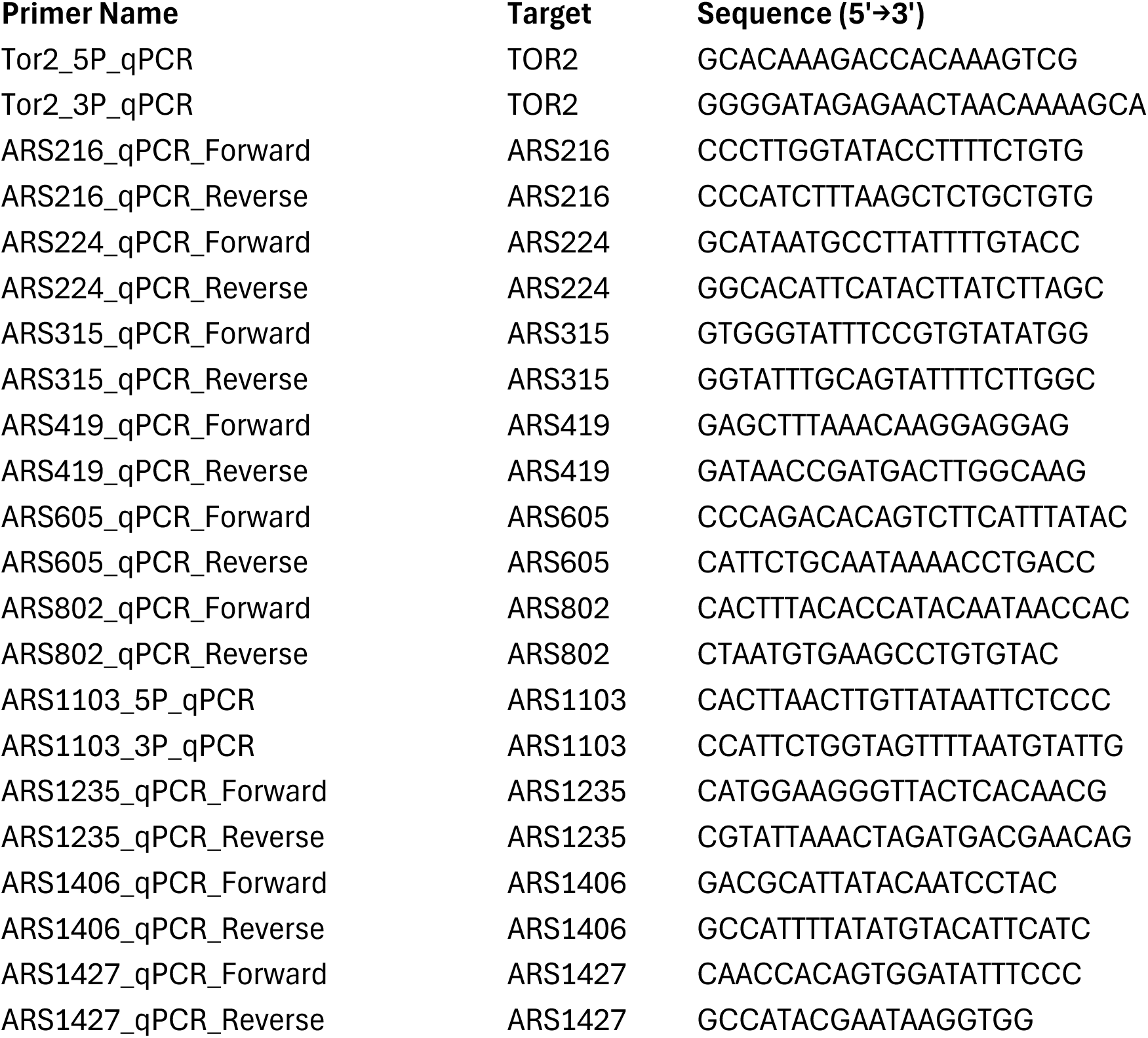
Primers used for qPCR.

## Notes

### Competing Interest Statement

The authors have declared no competing interest.

